# Deciphering mechanisms of UV filter- and temperature-induced bleaching in the coral *Acropora tenuis*, using ecotoxicogenomics

**DOI:** 10.1101/2023.11.05.564681

**Authors:** Sakiko Nishioka, Kaede Miyata, Yasuaki Inoue, Kako Aoyama, Yuki Yoshioka, Natsuko Miura, Masayuki Yamane, Hiroshi Honda, Toshiyuki Takagi

## Abstract

Coral reefs are at risk of bleaching due to various environmental and anthropogenic stressors such as global warming and chemical pollutants. However, there is little understanding of stressor-specific mechanisms that cause coral bleaching. Therefore, conducting accurate ecotoxicological risk assessments and deciphering modes of action of potentially deleterious ultraviolet (UV) filters (sunscreen compounds) are crucial issues. In this study, we evaluated the toxicity and bleaching effect of benzophenone-3 (BP-3), which is widely used in sunscreen products, on the reef-building coral *Acropora tenuis*. Furthermore, to understand differences in UV filter- and temperature-induced bleaching, a comparative ecotoxicogenomic approach using RNA-seq was integrated into a toxicity test to clarify differences in gene expression changes induced by BP-3 and heat stress (31°C). The lethal concentration 50% (LC50) was calculated as 3.9 mg/L, indicating that emission level of BP-3 was properly controlled based on the risk assessment. Differentially expressed genes related to oxidative stress and extracellular matrix organization were involved in coral responses to both BP-3 and heat stress, but their patterns differed. Whereas immune and heat-shock responses were activated in response to heat stress, activation of a drug metabolism pathway and several signal transduction pathways were identified in BP-3 treatment groups. Our study enhances understanding of stress responses in corals induced by UV filters and thermal stress. Using potential gene markers identified in this study for eco-epidemiological surveys of stressed corals, we urgently need to develop effective countermeasures.

## 1 Introduction

Preservation of marine biodiversity is a global concern to achieve nature-positive goals (Convention on Biological Diversity, 2022). Coral reef ecosystems are among the most productive and biodiverse on the planet. Although they cover less than 0.1% of the ocean floor, they provide food and shelter for approximately 25% of all marine organisms (Moeller et al., 2021; Spalding and Grenfell, 1997). They are not only ecologically crucial, but also economically important for food resources, tourism, and coastal protection (Souter and Lindén, 2000). Coral reef ecosystems are based on symbiotic relationships between host coral animals and endosymbiotic algae (Symbiodiniaceae). However, heat stress associated with global climate change induces a breakdown of symbiosis, i.e., coral bleaching (Sully et al., 2019). Prolonged bleaching leads to coral mortality, adversely impacting ecosystem services provided by coral reefs (Manzello et al., 2019).

In addition to heat stress, toxic effects of chemical pollutants in coral reefs are also a serious problem. Among them, ultraviolet (UV) filters, the active ingredients in sunscreens, may adversely affect corals and have attracted scientific, public, and regulatory attention (Bill SB 2571; State of Hawaii Senate, 2018; Sirois, 2019). Originally, this concern was raised by Danovaro *et al*. (2008), who suggested that organic UV filters cause coral bleaching at low concentrations through viral infections of Symbiodiniaceae. Subsequently, Downs *et al*. (2016) reported that an extremely high concentration of oxybenzone (benzophenone-3; BP-3) detected near the US Virgin Islands exceeded the lethal concentration 50% (LC50) for *Stylophora pistillata* larvae. Hence, the toxicological effects of UV filters, including BP-3, on corals were vigorously investigated using coral larvae and adult corals (He et al., 2019; Wijgerde et al., 2020; Conway et al., 2021). Consequently, a recent review suggested that environmental concentrations of UV filters are unlikely to exceed the toxicologically effective concentration for most corals (Mitchelmore et al., 2021).

However, the National Academy of Sciences of the USA identified an information gap regarding effects of UV filters on corals and recommended further research to protect both the environment and human health (National Academies of Sciences, Engineering, and Medicine et al., 2022). Despite increasing numbers of toxicity reports, the extent to which UV filters pose a risk to coral health remains unknown. One of the main reasons for this is that toxicity values vary widely among reports, which makes it difficult to accurately estimate toxicity. There are no guidelines for conducting coral toxicity tests. Thus, it is difficult to compare experimental conditions such as life cycle stages, test solution volumes, and exposure periods. For example, Downs *et al*. (2016) exposed coral larvae to BP-3 for 24 h, whereas He *et al*. (2019) exposed coral larvae for 14 d and adults for 7 d, and Danovaro *et al*. (2008) did not report exposure duration. Moreover, although some studies have measured and found a marked decrease of UV filter concentrations (He et al., 2019; Wijgerde et al., 2020), nominal concentrations were used to derive toxicity values.

Previous studies have not distinguished effects of UV filters and other environmental factors, especially heat stress. Although transcriptome analyses have been performed to investigate temperature-induced bleaching, and several potential genetic markers for heat stress have been reported (Cziesielski et al., 2019; Louis et al., 2017), no comprehensive analysis has investigated mechanisms underlying UV filter toxicity in corals. It is not known whether UV filters actually affect coral reefs, but genetic markers that can discriminate between UV filter- and temperature-induced bleaching may help identify possible threats in target coastal regions based on molecular eco-epidemiological studies.

The aim of this study was to understand the impact of BP-3 on corals under appropriate test conditions and to understand differences in BP-3- and temperature-induced bleaching, using comparative transcriptome analysis. We used the reef-building coral, *Acropora tenuis*, for BP-3 toxicity tests optimized with reference to OECD test guidelines. Furthermore, both BP-3 and heat stress groups were examined in a comparative transcriptome analysis using RNA-seq, based on the hypothesis that corals exposed to BP-3 respond differently than corals exposed to heat stress. This study provides mechanistic information on UV filter-induced toxicity from a gene expression perspective.

## 2 Material and Methods

### 2.1 Chemicals and preparation of sample solution

Oxybenzone (benzophenone-3; BP-3; 2-hydroxy-4-methoxybenzophenone) (CAS# 131-57-7, > 99.0%; FUJIFILM Wako Pure Chemical Corporation, Osaka, Japan) was dissolved in dimethyl sulfoxide (DMSO) (CAS# 67-68-5, > 99.0%; FUJIFILM Wako Pure Chemical Corporation, Osaka, Japan) at 100 g/L. Deuterated BP-3 (BP-3-d_5_, 2-Hydroxy-4-methoxybenzophenone-(phenyl-d_5_); CAS# 1219798-54-5, > 98%; Supelco Analytical) was used as the internal standard for liquid chromatography-triple quadrupole mass spectrometry quantification. All chemicals used were of analytical grade.

BP-3 dissolved in DMSO was added to artificial seawater at a nominal concentration of 10 mg/L, and the solution was stirred for 24 h with a magnetic stirrer at room temperature and then filtered through a 0.2 µm mixed cellulose ester membrane filter (Advantec Toyo, Tokyo, Japan) (OECD, 2019a). The filtered BP-3 solution was then diluted using filtered artificial seawater, and the final concentration of DMSO in each BP-3 solution was adjusted to 0.1 mL/L DMSO, following the maximum levels recommended by the OECD (OECD, 2019b). BP-3 solutions were prepared using only glassware to minimize adsorption.

### 2.2 Chemical analysis

To determine the actual BP-3 concentration during exposure, 100 mL of solution (n = 1) were collected from each test vessel and stored at -20°C until analysis. Samples were extracted by solid-phase extraction using Sep-Pak Plus tC18 cartridges with a polypropylene syringe, preconditioned with 10 mL of acetonitrile and 10 mL of distilled water. Samples (20 mL) were spiked with 0.20 mL BP-3-d_5_. After samples were introduced, cartridges were washed with 5.0 mL deionized water. Acetonitrile (8.0 mL) was added to the cartridge to elute BP-3 and was concentrated to 10 mL using deionized water. For highly concentrated samples, diluted samples were introduced into the cartridges. Liquid chromatography-mass spectrometry analysis was performed using the Agilent 1200 SL LC system coupled to an Agilent 6460 triple-quadrupole mass spectrometer (Agilent Technologies, Santa Clara, CA, USA). A 5.0-µL extracted sample was injected into a Mightysil RP-18 GP column (150 × 2.0 mm, 3.0 um; Kanto Chemical, Tokyo, Japan) at 40°C. The mobile phase was acetonitrile: deionized water (25:75), and the flow rate was held at 0.20 mL/min. Quantification was performed in electrospray ionization positive mode using precursor ions 229 and 234 for BP-3 and BP-3-d_5_, respectively, and product ion 151. The limit of quantification (LOQ), instrument quantification limit, and recovery rates of 10 µg/L BP-3 were 0.50 µg/L, 0.17 µg/L, and 99.2–102%, respectively.

### 2.3 Culture of coral Acropora tenuis

Corals were collected and cultured as previously described (Takagi et al., 2023). Briefly, six colonies of *A. tenuis* were collected from Sesoko Island, Okinawa, on December 7, 2020 and transported to the laboratory at the University of Tokyo (Kashiwa, Chiba Prefecture). Corals were kept at 26°C in an aquarium containing 600 L of artificial seawater prepared with Viesalt (Marine Tech, Tokyo, Japan). After one week of acclimation to the aquarium, apical branches approximately 3 cm in length were cut from each colony and mounted in a vertical position on ceramic coral frag plugs using Quick Gel (DELPHIS, Hyogo, Japan). All fragments were acclimated in the same aquarium for at least 8 weeks after fragmentation. Permits for coral collection were obtained from the Okinawa Prefectural Government (permit no. 2-46) for research use.

### 2.4 Dose range finding study

To assess the stability of BP-3 and to determine the appropriate exposure concentration range, a dose-finding study was performed by exposing fragments to four concentrations of BP-3 at a common ratio of 10 for 96 h (six fragments per group) in a static system. A blank control (BC) group and a solvent control (SC) group with 0.1 mL/L DMSO were also prepared. Test solutions were completely renewed only at the start of exposure. Other experimental designs were identical to those described in 2.5. During the 96-h exposure period, fragment mortality was recorded every 24 h.

### 2.5 Experimental design

Six fragments from different colonies were placed in each test vessel (n = 6) and acclimated for six days immediately before the start of the test. The fragments were exposed to five concentrations of BP-3 at a common ratio of 2.0 for 96 h, using a semi-static method (OECD, 2019b). Test solutions were completely renewed at the start of exposure and at 48 h, to maintain water quality and BP-3 concentration. The common ratio and renewal interval were determined based on a dose range finding study. BC and SC (0.1 mL/L DMSO) groups were also prepared. To minimize BP-3 adsorption, exposure was carried out in 40 × 26 × 30 cm glass vessels filled with 20 L of artificial seawater and glass equipment (Figure S1). Two glass tubes were used for aeration in each vessel to maintain dissolved oxygen (DO) levels and to provide constant random water flow. Glass rods were tied with polytetrafluoroethylene-coated yarn (Flon Industry Co., Ltd., Tokyo, Japan) to create a pedestal for coral fragments and were placed in the vessels. The vessels were kept in water baths maintained at 26 ± 1°C using heaters (Protect Heater R-300W plus SEAPALEX V-1000; NISSO, Kanagawa, Japan) and Voyager Nano stream pumps (SICCE, Italy). Vessels were equipped with LED lamps to provide the same light conditions as those of the aquarium.

To maintain water quality, salinity and water temperature were measured twice daily in each test vessel using a Marine Salinity Tester (HI98319; Hanna Instruments, Woonsocket, RI, USA). Moreover, a HOBO® Pendant light and temperature logger (UA-002-64; Onset Computer Corporation, Bourne, MA, USA) was placed in the BC vessel, and the temperature was recorded every 5 min throughout the experiment (Figure S2). Salinity was measured before and after addition of an appropriate amount of RO water to the vessels to compensate for water loss due to evaporation. DO levels (mg/L) and pH were measured daily using a Hanna Instruments® Portable Galvanic Dissolved Oxygen Meter (HI9147).

### 2.6 Biological analysis

During the 96-h exposure period, fragment mortality was recorded every 24 h. Fragments were photographed with a digital camera (Tough TG-5; Olympus, Tokyo, Japan). Light sensitivity (ISO 125), shutter speed (1/400 s), and aperture (f2.8) were held constant during the experiment. The distance between a fragment and the digital camera was fixed at 15 cm, and the Coral Color Reference Card (CCRC) (Siebeck et al., 2006) and color-matching sticker (CASMATCH, Bear Medic Corporation, Ibaraki, Japan) were included in the photographs. The CCRC was developed to record changes in coral color, providing a simple, inexpensive means of assessing the extent of coral bleaching in the field (Siebeck et al., 2006) and has been used in toxicity studies (Hirayama et al., 2017). Differences greater than two color scores of the CCRC reflect bleaching of the coral (Siebeck et al., 2006). All colors can be expressed as a combination of red, green, and blue brightness values, each of which is in the range of 0 (no brightness) to 255 (maximum brightness) in red-green-blue (RGB) color space. A black pixel has an RGB value of (0, 0, 0), and a white pixel has a value of (255, 255, 255). Adobe Photoshop CC^TM^ v19.1.6 (Adobe Systems, San Jose, CA, USA) was used for digital image processing and color quantification. First, photographs were calibrated using CASMATCH, according to the instructions. Next, the image area corresponding to the coral fragment was selected using the quick selection tool, and the RGB value was determined using the averaging tool. The square root of the sum of squares of the RGB values for each fragment was calculated. Those for the CCRC (D1–D6) included in the photographs of the BC group were also calculated, and standard curves were created for each imaging date. Fragment colors were quantified by converting RGB values to color scores using calibration curves.

The photosynthetic yield of photosystem II (PSII) of endosymbiotic algae was measured using a Junior pulse-amplitude modulated (PAM) fluorometer (Heinz Walz GmbH, Effeltrich, Germany) and PAM software (WinControl-3 v 3.29). Chlorophyll fluorescence has been used to assess coral bleaching, because a reduction in F_v_/F_m_ is linked to damage in PSII (Hoegh-Guldberg and Jones, 1999). Fragments were kept in the dark for 30 min before measurement for dark adaptation, and the dark-adapted maximum quantum yield of PSII (F_v_/F_m_) was recorded at three points (top, middle, and bottom) of each fragment.

### 2.7 Ecotoxicological risk assessment of BP-3

The predicted no-effect concentration (PNEC) is generally compared to environmental concentration such as the predicted environmental concentration (PEC) to determine a hazard quotient (HQ), and HQ > 1 indicates a potential cause for concern. In the present study, instead of PEC, the measured environmental concentration (MEC) was employed to assess the actual environmental risk, and we used the highest concentration (1.34 µg/L) reported in Okinawa in 2013 (Tashiro and Kameda, 2013) and other benchmark concentrations from a systematic review (Mitchelmore et al., 2021). Effective concentration 50% (EC50) was divided by an assessment factor of 1000 (European Chemicals Bureau, 2003), as it is more conservative than LC50 for deriving the PNEC.

### 2.8 Heat-stress experiment

A heat-stress experiment was simultaneously conducted to compare biological responses and gene expression between BP-3-exposed corals and heat-stressed corals. Six fragments from different colonies were placed in a glass vessel (n = 6) filled with 20 L of artificial seawater for the heat-stress treatment. The experimental design was the same as that for the BP-3 exposure treatment, and only the water temperature differed from that of the BC group (Figure S1C). After acclimation at 26°C for two days, water temperature was increased from 26 to 31°C at a rate of 1°C per day, and the 96-h heat-stress experiment was conducted on the same schedule as the BP-3 exposure experiment. A HOBO® Pendant temperature logger was also placed in the heat-stress group vessel (Figure S2). We previously conducted a long-term study using coral fragments derived from the same colonies as in this study, and confirmed that coral health was maintained even after 28 days in the control group (26°C), but bleaching was observed 3–4 weeks after the start of exposure at 31°C (Figure S3).

### 2.9 Sampling RNA extraction, RNA-seq library construction, and sequencing

At the end of the 96-h exposure period, all fragments from each vessel were removed from the ceramic coral frag plugs, and 5 mm of the growing tips were removed using pliers. Fragments were stored in RNAlater (Invitrogen, Carlsbad, CA, USA) at 4°C overnight and then at -80°C until use.

Fragments were ground to fine powder using an iron mortar and pestle and stored at -80°C. Total RNA was extracted from powdered samples using an RNeasy Plant Mini Kit (QIAGEN, Hilden, Germany) with the recommended DNase digestion. After extraction, the quantity of total RNA was determined using Synergy LX (BioTek, Winooski, VT, USA) and QuantiFluor RNA System (Promega, Madison, WI, USA). Total RNA quality was determined using a 5200 Fragment Analyzer System (Agilent Technologies, Santa Clara, CA, USA) and an Agilent HS RNA Kit (Agilent Technologies). RNA samples with an RNA Quality Number of ≥ 6.0 were used in our subsequent analysis. RNA libraries were constructed using an MGIEasy RNA Directional Library Prep Set (MGI Tech Co., Ltd., Shenzhen, China) according to the manufacturer’s instructions. Concentrations of these libraries were measured using Synergy H1 (BioTek) and QuantiFluor dsDNA System (Promega), and quality was checked using a Fragment Analyzer (Agilent Technologies) and dsDNA 915 reagent kit (Agilent Technologies). Circular DNA was prepared using constructed libraries and an MGIEasy circularization kit (MGI Tech Co., Ltd.), according to the manufacturer’s instructions. DNA nanoballs (DNBs) were prepared using the DNBSEQ-G400 RS High-throughput Sequencing Kit (MGI Tech Co., Ltd.), according to the manufacturer’s instructions. DNBs were sequenced using a DNBSEQ-G400 (MGI Tech Co., Ltd.) under 2 × 200-bp conditions.

### 2.10 Differentially expressed gene analysis

Low-quality reads (quality score < 20 and length < 100 bp) and DNBSEQ sequence adaptors (Read1: 5’-AAGTCGGAGGCCAAGCGGTCTTAGGAAGACAA-3’; Read2: 5’-AAGTCGGATCGTAGCCATGTCGTTCTGTGAGCCAAGGAGTTG-3’) were trimmed using Cutadapt v4.0 (Martin, 2011). Cleaned reads were mapped to the *A. tenuis* gene models (mRNA) (Shinzato et al., 2021), downloaded from the genome browser of the Okinawa Institute of Science and Technology Marine Genomics Unit (https://marinegenomics.oist.jp), using Salmon v1.8.0 (Patro et al., 2017) with default settings. Mapped read counts were normalized using the trimmed mean of M values (TMM) method and converted to counts per million using EdgeR (version 3.32.1) (Robinson et al., 2010) in R (version 4.0.3). TMM-normalized mapped read counts in treatment groups were compared pairwise with control samples to identify differentially expressed genes (DEGs). During pairwise comparison, genes with low expression were filtered out using the “filterByExpr” function in EdgeR. *P*-values were adjusted using the Benjamini–Hochberg method in EdgeR. Genes were considered DEGs if the gene expression level was significantly different (false discovery rate (FDR) < 0.05, and |log 2 fold change (log2FC)| > 0.58) from those of control samples. Gene annotation was performed by searching the Swiss-Prot eukaryotic protein database (downloaded on November 25, 2022) using BLASTX with an e-value cutoff of *E* = 10^-5^. When the same gene symbol was assigned, the serial number was appended after an underscore. Numbers of DEGs and overlaps were visualized using the VennDiagram package (version 1.7.3) and expression levels of each DEG were visualized using ggplot2 (version. 3.4.1) in R (version 4.2.2).

### 2.11 PCA analysis, enrichment analysis and pathway analysis

Principal component analysis (PCA) was conducted on the DEGs (FDR < 0.05 and |log2FC| > 0.58) of each group and visualized using the factoextra package (version 1.0.7) in R. Principal components (PCs) that were associated with discrimination of the BP-3 treatment and heat-stress groups were explored, DEGs with cos2 values greater than 0.8 or particularly highly upregulated or downregulated DEGs (top 20 based on log2FC in each gene set) were extracted, and groups of genes with characteristic functions were manually curated. The cos2 value represents the quality of the representation of the variable, with a high cos2 indicating that the variable is well represented on the PCs.

To investigate the biological classification of DEGs, Gene Ontology (GO) biological process (BP) and Reactome pathway enrichment analyses were performed for each gene set (only for annotated genes) using Database for Annotation, Visualization, and Integrated Discovery (DAVID) software (https://david.ncifcrf.gov/). Significantly enriched terms (FDR < 0.1) in either gene set were extracted and depicted as a heatmap using the pheatmap package (version 1.0.12) in R.

Data were analyzed through the use of IPA (QIAGEN Inc., https://www.qiagenbioinformatics.com/products/ingenuity-pathway-analysis) to evaluate functional networks among DEGs. Only direct biological relationships were analyzed and genes with at least one or more relationships to other genes were extracted and visualized. Genes are represented as nodes, and biological relationship between nodes are represented as edges (lines). Representative canonical pathways were determined based on numbers of genes mapped to a pathway.

### 2.12 Statistical analysis

96-h LC50 and EC50 values were estimated based on measured concentrations using Ecotox-Statics (version 3.01.01) software (the Japanese Society of Environmental Toxicology). One-way ANOVA, followed by the Tukey–Kramer test, was employed to evaluate statistical differences in body color and F_v_/F_m_ values for multiple comparisons among groups. This analysis was performed to determine differences between measurements at the beginning and end of the treatment. Furthermore, paired *t*-tests were employed to analyze statistical differences at time point 0, and *p*-values were adjusted using the Holm method. Paired *t*-tests were performed using Microsoft Excel, and Tukey–Kramer tests were performed using the multcomp package in R (version 4.2.2) and R Studio (version 2022.07.2). Differences were considered statistically significant at *p* < 0.05.

## 3 Results

### 3.1 Dose finding test

Concentrations at the beginning of exposure were 6.5, 0.69, 0.068, and 0.0058 mg/L (Table S1). Mortality rates of 100% were observed only at the highest concentration, and no mortality was observed at lower concentrations or in control groups, and so a common ratio of 2 was applied for the definitive test. At the end of the BP-3 exposure period, concentrations of BP-3 were 6.1 (at the 24-h time point), 0.18, 0.055 mg/L, and below the LOQ, respectively. Due to its relatively high lipophilicity (LogKow = 3.45; ECHA. 2011), a certain amount of BP-3 can adsorb to equipment surfaces or to coral mucus, reducing measured concentrations (Conway et al., 2021; He et al., 2019; Wijgerde et al., 2020).

### 3.2 Water quality and maintenance of BP-3 concentration

Water quality parameters (temperature, salinity, pH, and DO) were stable throughout the experiment and similar among vessels (Table S2). Water temperatures of the BC and heat-stress groups were maintained in the range of 25.7–26.7°C and 30.9–31.6°C, respectively (Figure S2).

No BP-3 was detected in the BC or SC groups throughout the experiment, and the mean concentrations of BP-3 in the new test solutions were 5.9, 3.0, 1.7, 0.83, and 0.41 mg/L (Table 1). The maximum concentration was only slightly lower than the reported solubility in water (ECHA, 2011), suggesting that 5.9 mg/L was the solubility limit in the test medium under these conditions, and corals were only exposed to dissolved BP-3 fractions. BP-3 concentrations remained above 80% from the initial concentrations throughout the test. Time-weighted mean (TWM) concentrations were 5.6, 2.7, 1.5, 0.77, and 0.38, and retention rates of TWM concentrations from mean initial concentrations were 95.7%, 90.8%, 93.1%, 93.6%, and 93.8%, respectively, indicating that BP-3 did not degrade or adsorb markedly under these experimental conditions. Although the deviation from the initial concentrations was less than 20%, we used the measured concentrations of BP-3 recommended in the OECD test guidelines to understand the toxicity of BP-3 to *A. tenuis* (OECD, 2019b) precisely. Based on measured concentrations, BP-3 treated groups were named BP-3_5.6, BP-3_2.7, BP-3_1.5, BP-3_0.77, and BP-3_0.38.

**Table 1.**
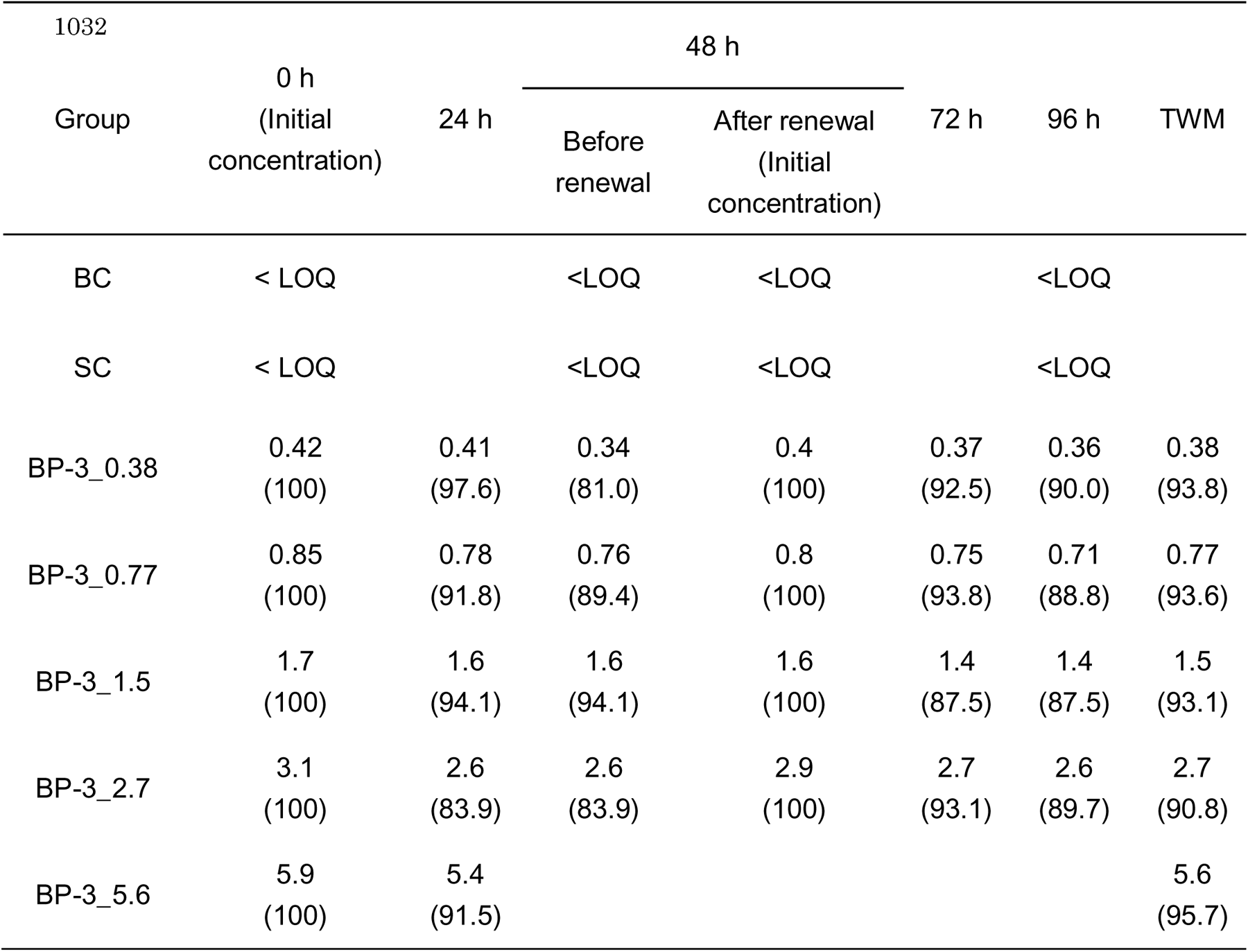
Measured concentrations of BP-3 (mg/L) in test solutions (n = 1). The number after the underscore in each group name represents the time-weighted mean (TWM) concentration of BP-3. The retention rate at each time point compared with the initial concentration is shown in parentheses (%). Retention rates of TWM values were calculated by dividing TWM concentrations by mean initial concentrations. At 48 h, test solutions were analyzed before and after renewal. LOQ: limit of quantification, BC: blank control, SC: solvent control.

### 3.3 General Observations

Major observations of BP-3-exposed and heat-stressed corals are summarized in Table 2. Photographs of fragments at the beginning and end of treatment are shown in Figure 1A. Fragment mortality in control vessels was 0%, fulfilling the validation criteria of OECD test guidelines (OECD, 2019b). Rapid tissue lysis and 100% mortality were only observed in group BP-3_5.6 within 12 h and dead fragments had exposed white skeletons (Figure 1A); therefore, the exposure experiment was terminated at 24 h only for this group. At the second highest BP-3 concentration (BP-3_2.7), all fragments had severely retracted polyps, and partial bleaching was observed in some areas, such as specific polyps and edges of the fragments.

**Figure 1.**
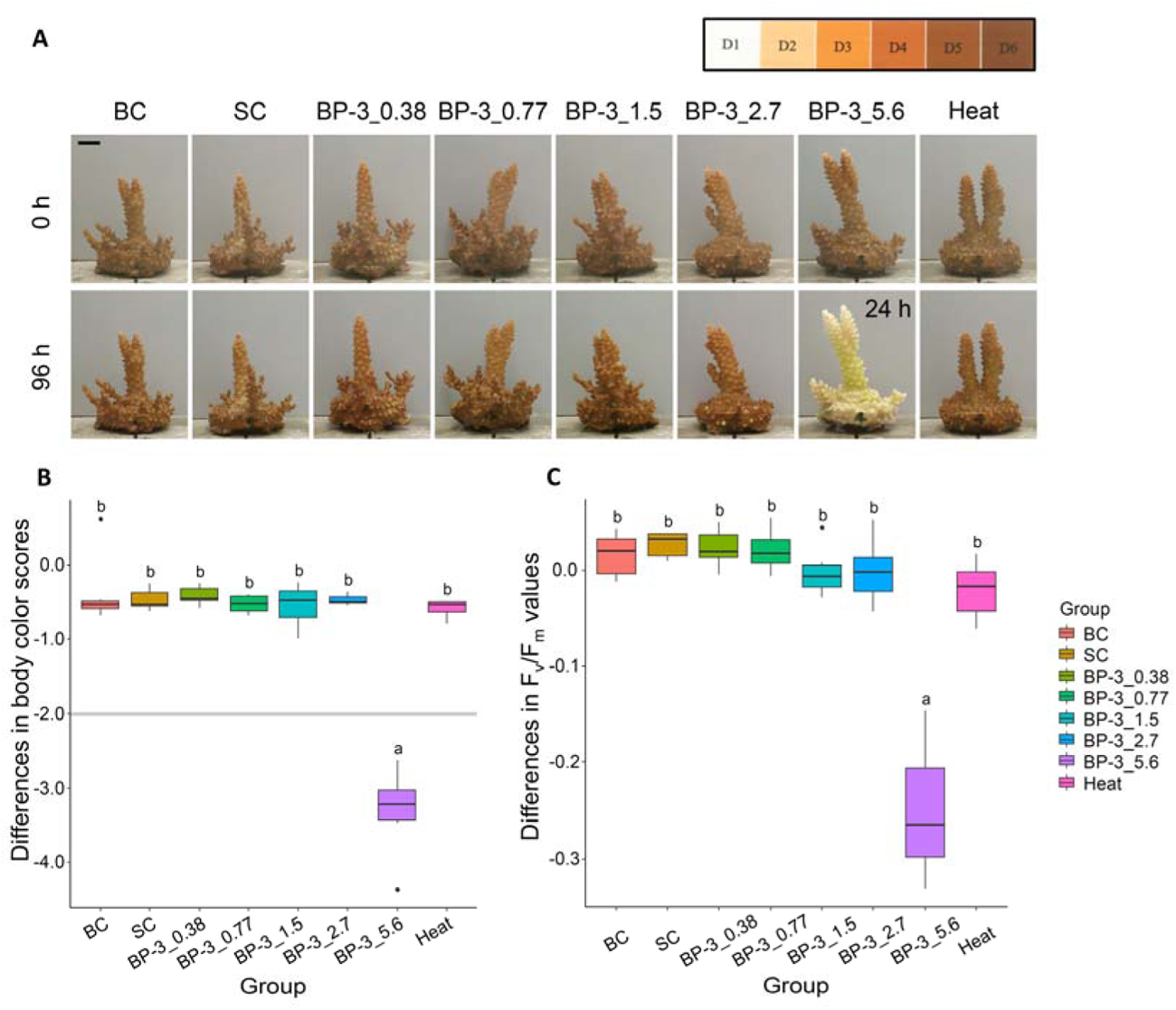
Comparisons of body color scores and photosynthetic activities. (A) Photographs of *Acropora tenuis* exposed to BP-3 or heat stress for 96 h (except for BP-3_5.6, in which all fragments were dead at 24 h). Presented fragments were from the same colony. A color reference card (Siebeck et al., 2006) is shown in the upper right corner. Differences in (B) body color score and (C) F_v_/F_m_ values of *A. tenuis* between the start (0 h) and end (96 h, except for BP-3_5.6) of the treatment. The grey line indicates the bleaching criterion (Siebeck et al., 2006). Different letters indicate significant differences (Tukey–Kramer test). Bar = 1 cm, BC: blank control, SC: solvent control.

**Table 2.** Summary of biological endpoints of BP-3 and heat-stress exposure in the coral, *Acropora tenuis*. The number after the underscore in each group name represents the time-weighted mean (TWM) measured concentration of BP-3. Color scores and F_v_/F_m_ values are presented as the average ± standard deviation for each test vessel. TWM: time-weighted mean, LOQ: limit of quantification, BC: blank control, SC: solvent control.

| Group | TWM<br>concentration<br>(mg/L) | Mortality<br>(%) | Observation | Color Score | | $F_v/F_m$ | |
| --- | --- | --- | --- | --- | --- | --- | --- |
|  |  |  |  | Start | End | Start | End |
| BC | < LOQ | 0 | Not<br>changed | 6.3<br>±0.46 | 6.0<br>±0.74 | 0.61<br>±0.023 | 0.63<br>±0.011 |
| SC | < LOQ | 0 | Not<br>changed | 6.3<br>±0.45 | 5.8<br>±0.45 | 0.60<br>±0.024 | 0.63<br>±0.023 |
| BP-3_0.38 | 0.38 | 0 | Not<br>changed | 6.6<br>±0.45 | 6.2<br>±0.44 | 0.61<br>±0.016 | 0.63<br>±0.020 |
| BP-3_0.77 | 0.77 | 0 | Not<br>changed | 6.6<br>±0.37 | 6.1<br>±0.40 | 0.61<br>±0.034 | 0.63<br>±0.022 |
| BP-3_1.5 | 1.5 | 0 | Not<br>changed | 6.6<br>±0.37 | 6.0<br>±0.40 | 0.62<br>±0.031 | 0.64<br>±0.025 |
| BP-3_2.7 | 2.7 | 0 | Partially<br>bleached<br>Polyp<br>retraction | 6.4<br>±0.36 | 6.0<br>±0.40 | 0.61<br>±0.021 | 0.61<br>±0.034 |
| BP-3_5.6 | 5.6 | 100<br>within<br>12 h | Tissue lysis<br>within 12 h | 6.5<br>±0.42 | 3.2<br>±0.39 | 0.61<br>±0.024 | 0.36<br>±0.055 |
| Heat | - | 0 | Not<br>changed | 6.7<br>±0.52 | 6.1<br>±0.55 | 0.61<br>±0.028 | 0.60<br>±0.028 |

### 3.4 Effect on body color and photosynthetic activity

Differences in body color score of *A. tenuis* at the beginning and end of treatment were compared (Figure 1B and Table 2). During the treatment period, color scores decreased slightly, but significantly in all groups except the BC group (paired *t*-test). However, the mean change for all groups, except for the highest BP-3 concentration group (BP-3_5.6), was not significant (Tukey–Kramer test) and changes in color scores (0.3–0.6) did not exceed the bleaching criterion (Siebeck et al., 2006). The mean change in BP-3_5.6 group was 3.3 at 24 h, which exceeded the bleaching criterion, with statistically significant differences (*p* < 0.05).

Body color in the heat-stress group showed a slight decreasing trend, but this was not significant. A preliminary study has confirmed that exposure to 31°C for 3–4 weeks causes bleaching (Figure S3). Since bleaching is caused by cumulative heat stress, this is an appropriate time period to capture changes that could predict temperature-induced bleaching.

F_v_/F_m_ values of all groups were approximately 0.61 (Table 2) at the beginning of the treatment; however, the mean F_v_/F_m_ value of the highest BP-3 concentration group (BP-3_5.6) decreased significantly (*p* < 0.05) at 24 h (Figure 1C). At lower BP-3 concentrations (0.38, 0.77, 1.5, and 2.7 mg/L), values remained constant, indicating that photosynthetic activity of endosymbiotic algae was maintained during the exposure period, which is consistent with the results of the body color analysis (Figure 1B).

### 3.5 Derivation of toxicological values

The LC50 was calculated as 3.9 mg/L based on quantitative evaluation results. Although no effects on mortality, body color score, or F_v_/F_m_ were found at 2.7 mg/L, severely retracted polyps and partial bleaching were observed. Thus, the no observed effect concentration (NOEC), lowest observed effect concentration (LOEC), and EC50 were 1.5 mg/L, 2.7 mg/L, and 2.0 mg/L, respectively.

### 3.6 Gene responses of coral to BP-3 and heat stress

Transcriptomes of *A. tenuis* under BP-3 exposure and heat stress were sequenced to compare its gene responses and to identify potential gene markers for distinguishing BP-3-and temperature-induced bleaching. Numbers of mapped reads in each sample ranged from 20,695,558 to 32,383,474, and mapping rates ranged from 41% to 53% (Table S3). Fragments exposed to 5.6 mg/L BP-3 were excluded from transcriptome analysis because they suffered 100% mortality and rapid tissue lysis within 12 h, and RNA extraction was not possible for this group. Comparison of the BC and SC groups was conducted first, and there were only two DEGs; therefore, BP-3-exposed groups were compared to the SC group, and the heat-stressed group was compared to the BC group because no solvent was added to the heat-stressed group.

In the BP-3_2.7 group, 2,369 DEGs were identified (Figure 2A), and 1,602 (67.6%) genes were annotated using the Swiss-Prot database. DEGs were also identified at lower BP-3 concentrations (28 genes in BP-3_0.38, 58 genes in BP-3_0.77, and 239 genes in BP-3_1.5), and 17 (60.7%), 35 (60.3%), and 155 (64.9%) genes were annotated, respectively. Numbers of DEGs increased dramatically with increasing BP-3 concentrations. To compare responses between BP-3 exposure and heat stress, DEGs at either BP-3 concentration were compared to DEGs in heat-stressed group (Figure 2B, Figure S4). A total of 1,878 DEGs were BP-3 specific, 339 DEGs were heat-stress specific, and 522 DEGs were common to both. 1,279 (68.1%), 240 (60.2%), and 345 (66.1%) genes were annotated in the three groups, respectively, using the Swiss-Prot database.

**Figure 2.**
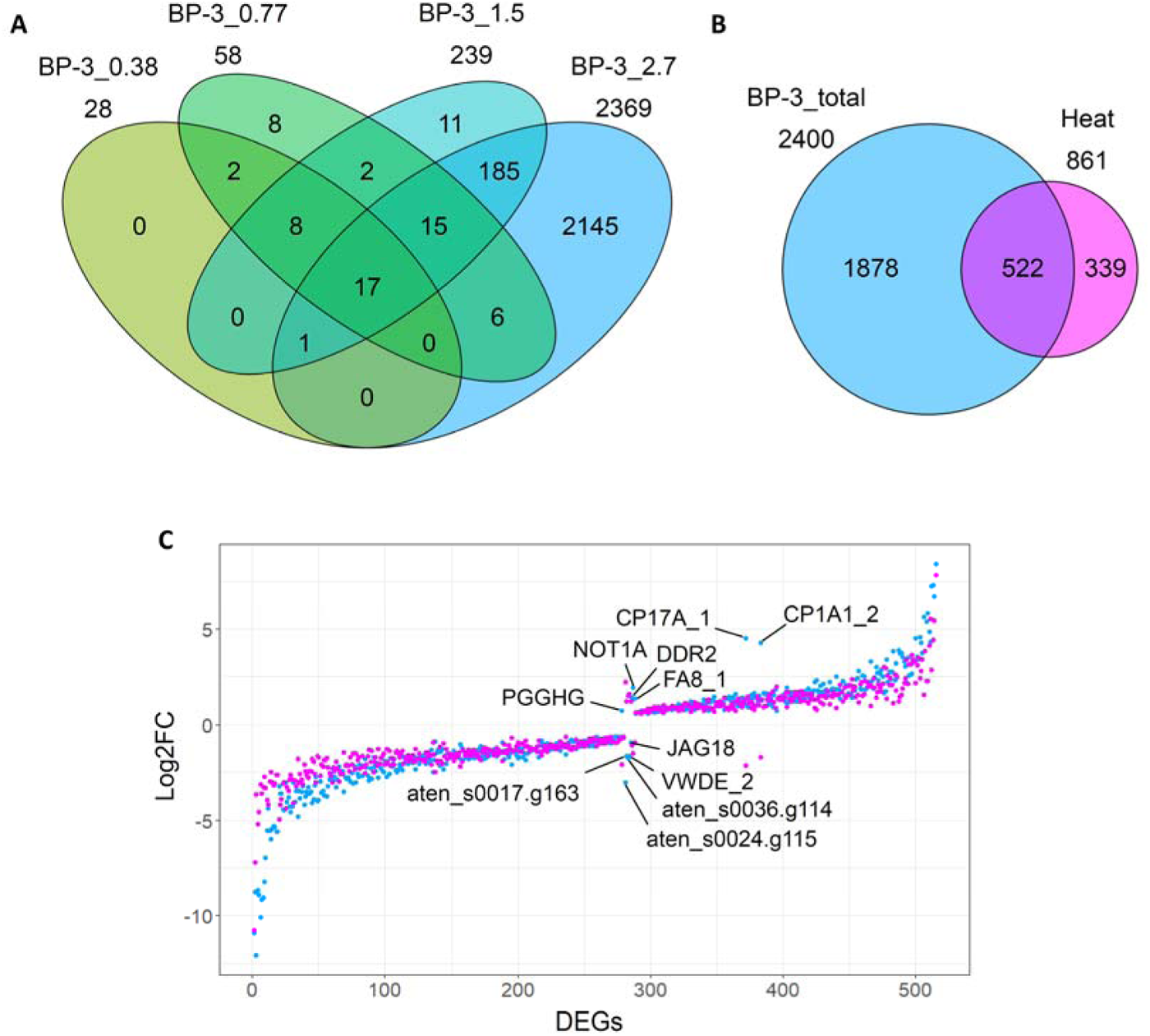
Comparison of differentially expressed genes (DEGs) in BP-3 and heat-stress exposure groups. (A) Venn diagram of numbers of DEGs in BP-3 concentration groups. (B) Venn diagram of DEGs that showed expression variation upon exposure to any concentration of BP-3 (sum assembly of DEGs shown in Figure 2A) and DEGs upon heat stress. (C) Log2FC plots of 522 DEGs shared between BP-3 and heat-stress exposure groups. For common DEGs with mean values of BP-3_2.7 and heat stress in ascending order, the vertical axis represents expression variation. Cyan-colored plots show expression variation (log2FC) in BP-3_2.7 compared with the solvent control (SC) group. Magenta-colored plots show expression variation (log2FC) in heat-stressed plots compared to the blank control (BC) group. Among common DEGs with differential expression variation patterns under the two stress sources, genes annotated in Swiss-Prot are described.

Among BP-3-specific DEGs, aten_s0046.g96 (*galaxin: GXN_3*) had the highest increase in gene expression level (Log2FC = 9.40 in BP-3_2.7), followed by aten_s0342.g22 (*peroxidasin homolog 2: PXDN2_1*) (Log2FC = 8.09 in BP-3_2.7) and aten_s0342.g21 (*PXDN_3*) (Log2FC = 7.53 in BP-3_2.7) (Table S4). Most genes for which expression varied under both BP-3 exposure and heat stress showed similar gene expression changes (Figure 2C). On the other hand, the following DEGs showed expression changes in the opposite direction: aten_s0001.g47 (*cytochrome P450 family 1 subfamily A member 1: CP1A1_2*), aten_s0012.g187 (*steroid 17 alpha-hydroxylase/17, 20-lyase: CP17A_1*), aten_s0035.g108 (*coagulation factor VIII: FA8_1*), aten_s0059.g52 (*palmitoleoyl-protein carboxylesterase notum1a: NOT1A*), aten_s0240.g12 (*discoidin domain receptor tyrosine kinase 2: DDR2*), aten_s0035.g8 (*jagged canonical Notch ligand 1b: JAG1B*), aten_s0211.g24 (*von Willebrand factor D and epidermal growth factor domains: VWDE_2*), aten_s0052.g91 (*protein-glucosylgalactosylhydroxylysine glucosidase: PGGHG*), aten_s0036.g114, aten_s0017.g163, and aten_s0024.g115.

### 3.7 PCA for identification of potential gene markers

To explore the variation in gene expression, we performed PCA for identified DEGs. As mentioned above, there were only two DEGs between BC and SC, and they were plotted at overlapping positions, confirming that effects of the solvent control (DMSO) were minor (Figure 3A). Six groups (BC, SC, BP-3_0.38, BP-3_0.77, BP-3_1.5, and BP-3_2.7) were plotted in a straight line, and distance from the control groups increased with increasing BP-3 concentration. In contrast, the heat-stress group was plotted at a large distance from the straight line described above. The first principal component (PC1) accounted for 43.4% of the variation, reflecting a change in the concentration-dependent gene expression profile following BP-3 exposure. Expression levels of PC1-related genes varied most remarkably in group BP-3_2.7. Subsequently, these genes were similarly variable in the heat stress and BP-3_1.5 groups, indicating that PC1-related genes are also associated with the common mechanism of BP-3 and temperature-induced responses. In contrast, the contribution rate of the second component (PC2) was 11.2% and remarkably discriminated the genetic expression profile of the heat-stress group from those of BP-3 exposed groups.

**Figure 3.**
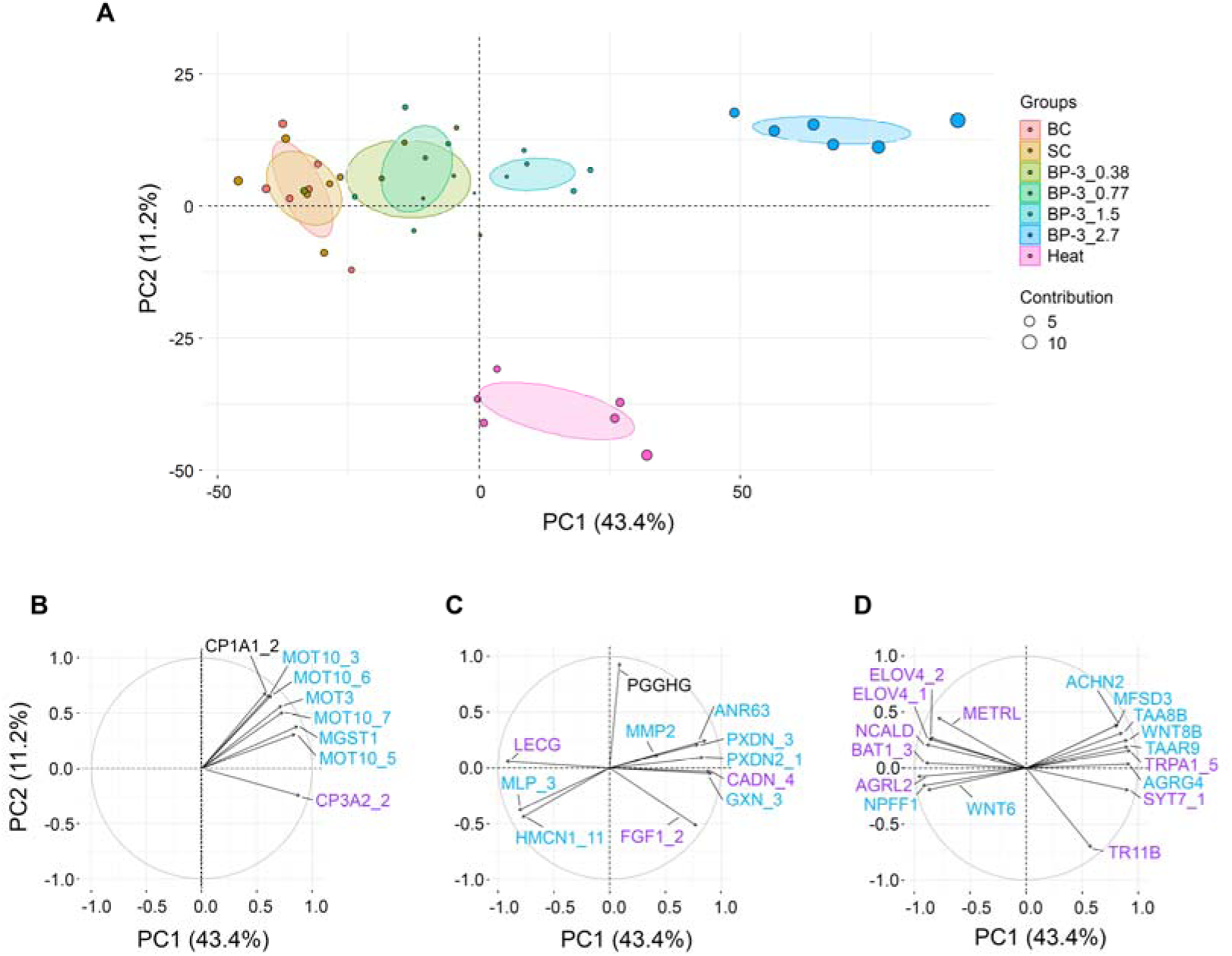
Separation of BP-3 and heat-stress exposure groups by principal component analysis (PCA) and associated genes. (A) PCA plot based on log2FC values of all differentially expressed genes (DEGs) that showed expression differences in control and treatment groups. Plots indicate each sample with 95% confidence intervals, and the size of plots indicates the contribution to the principal component of each sample. Contributions of the first (PC1) and second principal component (PC2) were 43.4% and 11.2%, respectively. BC: blank control, SC: solvent control, Heat: heat-stress. (B–D) Biplot extracted for (B) metabolism-related factors, (C) extracellular matrix (ECM) and cytoskeleton-related factors, and (D) signal transduction-related factors among the DEGs annotated with cos2 > 0.8 or with particularly highly differentially expressed DEGs, described in Table S4. cyan: BP-3-specific DEGs, purple: common DEGs, and black: opposite expression changes (Figure 2B, C).

Based on the cos2 value, DEGs were extracted to identify potential gene markers that distinguish BP-3-and temperature-induced bleaching. Of the annotated DEGs with cos2 > 0.8 or especially highly upregulated or downregulated DEGs described in Table S4, notable DEGs were classified into three categories: metabolism-related factors (Figure 3B); extracellular matrix (ECM) organization and cytoskeleton-related factors (Figure 3C); signal transduction-related factors (Figure 3D).

Metabolism-related factors included genes related to drug disposition such as metabolic enzymes and transporters: *CP1A1_2*, aten_s0087.g22 (*CP3A2_2*), aten_s0049.g117 (*microsomal glutathione S-transferase 1: MGST1*), aten_s0081.g13 (*monocarboxylate transporter 10: MOT10_3*), aten_s0034.g34 (*MOT10_5*), aten_s0027.g65 (*MOT10_6*), aten_s0326.g13 (*MOT10_7*), and aten_s0115.g39 (*MOT3*) (Figure 3B). ECM-related factors included *GXN_3*, aten_s0097.g15 (*coadhesin: CADN_4*), aten_s0055.g111 (*hemicentin 1: HMCN1_11*), aten_s0067.g146 (*ankyrin repeat domain-containing protein 63: ANR63*), aten_s0127.g24 (*mucin*-like *protein: MLP_3*), aten_s0046.g80 (*galactose-specific lectin nattectin: LECG*) (Figure 3C). Signal transduction-related factors included aten_s0085.g90 (*adhesion G protein-coupled receptor L2: AGRL2*), aten_s0117.g81 (*neuropeptide FF receptor 1: NPFF1*), aten_s0005.g5 (*neuronal acetylcholine receptor subunit_non-alpha-2: ACHN2*), aten_s0027.g41 (*neurocalcin-delta: NCALD*), aten_s0046.g59 (*synaptotagmin-7: SYT7_1*), aten_s0002.g104 (*transient receptor potential cation channel subfamily A member 1: TRPA1_5*), aten_s0350.g6 (*wingless-type MMTV integration site family, member 6: WNT6*), aten_s0336.g16 (*WNT8*B), aten_s0075.g72 (*tumor necrosis factor receptor superfamily member 11B: TR11B*), aten_s0006.g150 (*major facilitator superfamily domain-containing protein 3: MFSD3*), aten_s0150.g17 (*meteorin*-like *protein: METRL*), aten_s0139.g63 (*b(0,+)-type amino acid transporter 1: BAT1_3*), aten_s0068.g56 (*elongation of very long-chain fatty acids protein 4: ELOV4_1*), aten_s0068.g57 (*ELOV4_2*), aten_s0075.g100 (*trace amine-associated receptor 9: TAAR9*), aten_s0114.g18 (*adhesion G protein-coupled receptor G4: AGRG4*), and aten_s0038.g19 (*trace amine-associated receptor 8b: TAA8B*) (Figure 3D).

### 3.8 O and Reactome pathway enrichment analysis

To identify molecular mechanisms involved in the response to BP-3 exposure and heat stress, GO-BP and Reactome pathway enrichment analyses were conducted for annotated genes in each gene set in Figure S4 (Figure 4). Significantly enriched terms (FDR < 0.1) in each dataset are summarized in Table S5. BP-3-specific upregulated DEGs (Up-DEGs) were enriched in the G protein-coupled receptor (GPCR) signaling pathway (GO:0007186), extracellular matrix organization (GO:0030198), the fibroblast growth factor receptor signaling pathway (GO:0008543), and heart looping (GO:0001947). BP-3-specific downregulated DEGs (Down-DEGs) were enriched in chloride transmembrane transport (GO:1902476), neurological system processes (GO:0050877), regulation of presynaptic membrane potential (GO:0099505), chemical synaptic transmission (GO:0007268), homophilic cell adhesion via plasma membrane adhesion molecules (GO:0007156), excitatory postsynaptic potential (GO:0060079), regulation of membrane potential (GO:0042391), visual perception (GO:0007601), neuropeptide signaling pathway (GO:0007218), ion transmembrane transport (GO:0034220), defense response (GO:0006952), and angiogenesis (GO:0001525).

**Figure 4.**
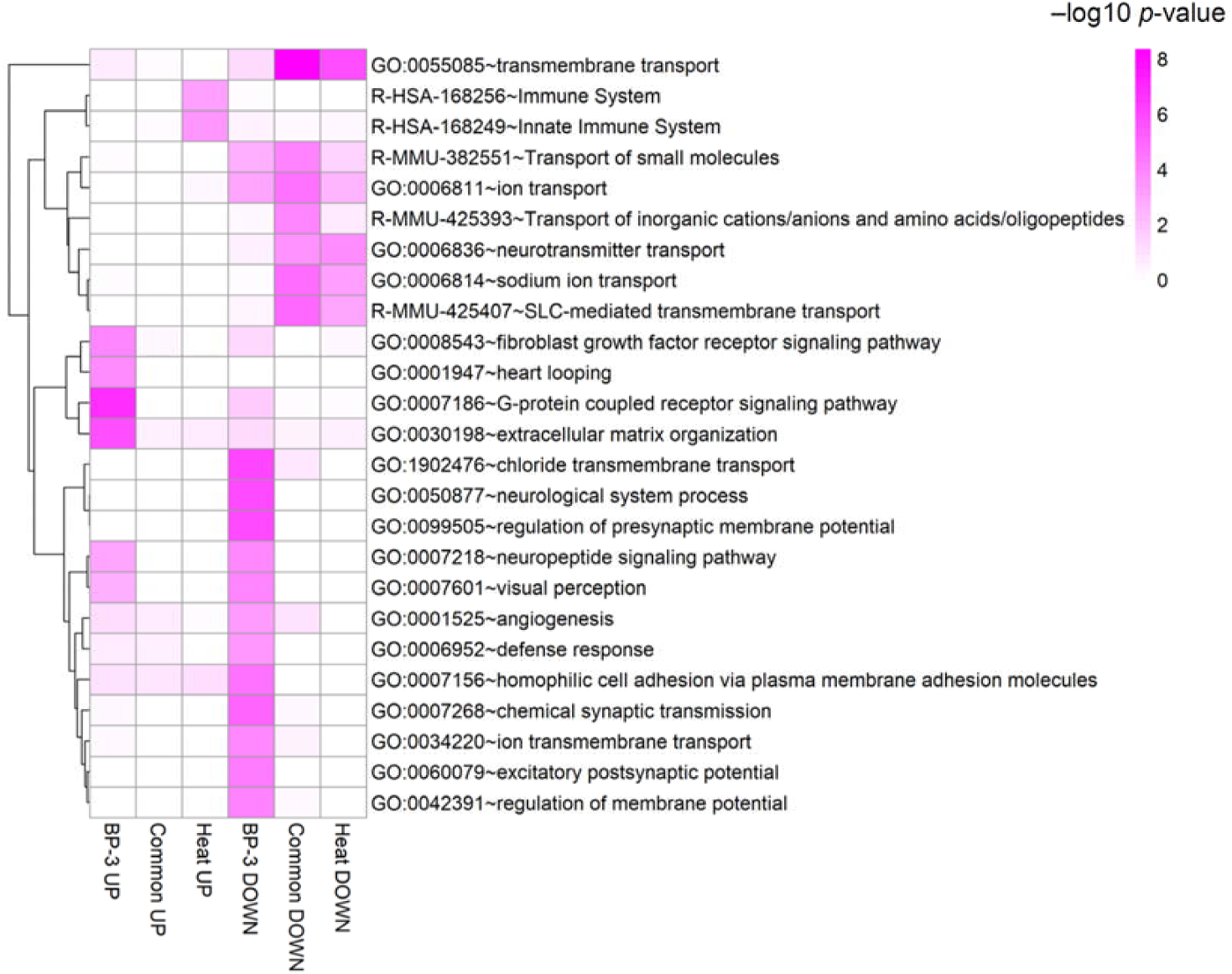
Enrichment analysis of differentially expressed genes (DEGs). Gene Ontology (GO) biological process (BP) terms and Reactome pathways with FDR < 0.1 in each group (BP-3-specific Up-DEGs and Down-DEGs, heat-specific Up-DEGs and Down-DEGs, and common Up-DEGs and Down-DEGs) in Figure S4. Each cell in the heatmap is colored based on the –log10 *p*-value. GO: gene ontology; BP: biological process; Up-DEGs: upregulated differentially expressed genes, Down-DEGs: downregulated differentially expressed genes. The darker the magenta, the higher the –log10 *p*-value. Terms starting with ‘GO:’ indicate BP and those starting with ‘R-’ indicate Reactome pathways.

Heat-specific Up-DEGs were enriched in the innate immune system (R-HSA-168249) and immune system (R-HSA-168256), whereas heat-specific Down-DEGs were enriched in transmembrane transport (GO:0055085) and neurotransmitter transport (GO:0006836). Common Down-DEGs were enriched in transmembrane transport (GO:0055085), SLC-mediated transmembrane transport (R-MMU-425407), sodium ion transport (GO:0006814), ion transport (GO:0006811), transport of small molecules (R-MMU-382551), transport of inorganic cations/anions and amino acids/oligopeptides (R-MMU-425393), and neurotransmitter transport (GO:0006836). The GO term “transmembrane transport” overlapped between heat-specific Down-DEGs and common Down-DEGs.

### 3.9 Gene network analysis

To understand networks based on interactions between DEGs and to estimate their functions, pathway analysis was conducted. The functional gene network analysis of BP-3-specific, heat-specific, and common Up-DEGs were conducted (Figure S4A). For BP-3-specific Up-DEGs, molecules associated with the top four canonical pathways (S100 family signaling pathway, axonal guidance signaling, myelination signaling pathway, and G protein-coupled receptor signaling pathway) are depicted in Figure 5A. Common Up-DEGs included aten_s0140.g17 (*mitogen-Activated Protein Kinase 14: MAPK14*), aten_s0207.g13 (*jun proto-oncogene: JUN*), aten_s0087.g22 (*CYP3A4*), and aten_s0316.g12 (*CYP3A 5*), which were associated with several canonical pathways such as NRF2-mediated oxidative stress response and xenobiotic metabolism signaling (Figure 5B). For heat-specific Up-DEGs, *heat shock proteins* (*HSPs*) aten_s0008.g125 (*HSPA5*) and aten_s0006.g70 (*HSP90B1*) were included in the network (Figure 5C), which is consistent with previous reports (Louis et al., 2017). They are associated with canonical pathways such as the NOD 1/2 signaling pathway, the role of PKR in interferon induction and antiviral response, and the immunogenic cell death signaling pathway.

**Figure 5.**
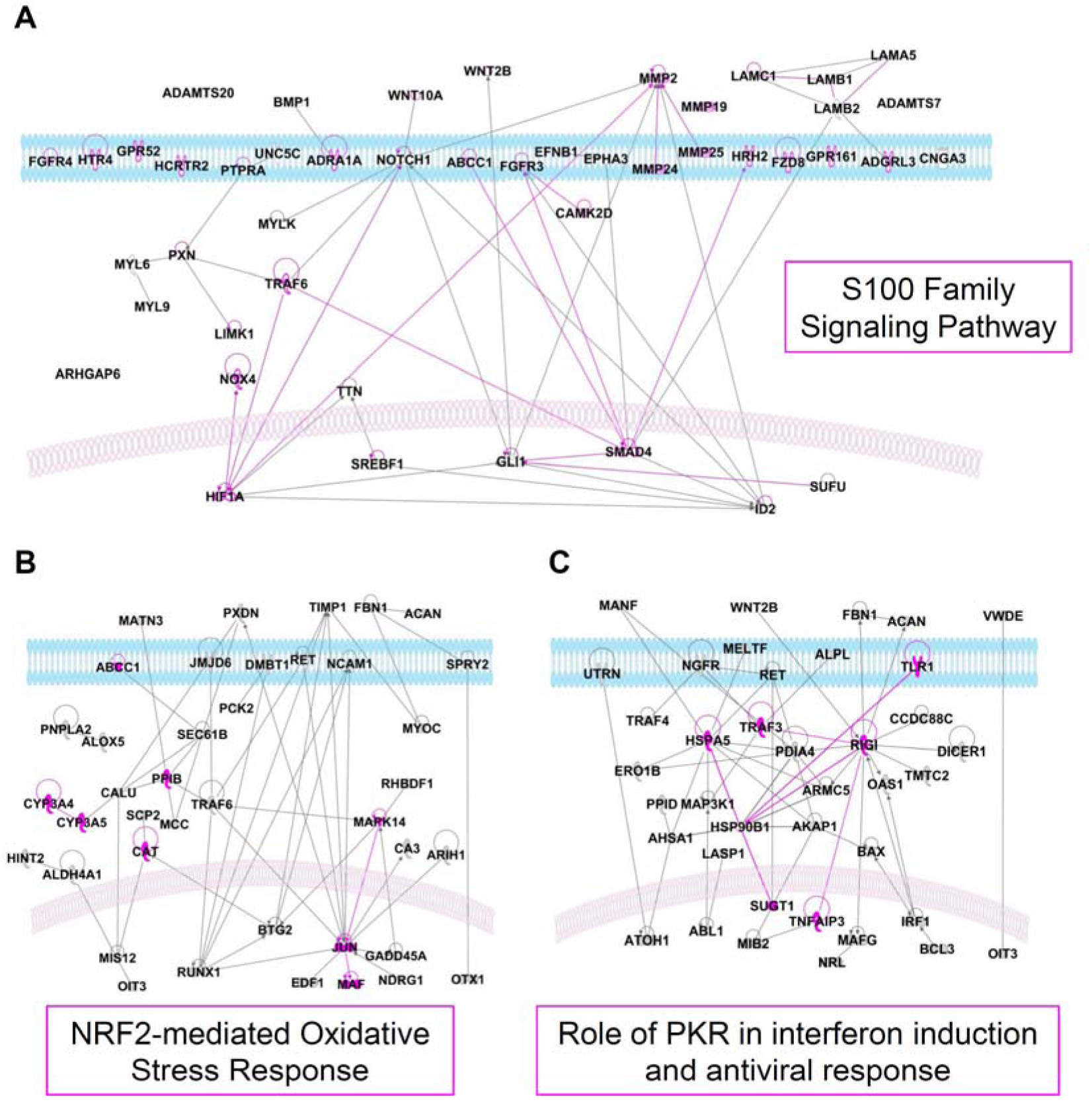
Gene network analysis of differentially expressed genes (DEGs). Ingenuity Pathway Analyses (IPAs) were conducted for (A) BP-3-specific upregulated DEGs (Up-DEGs), (B) common Up-DEGs and (C) heat-specific Up-DEGs. For BP-3-specific Up-DEGs, molecules associated with the top four canonical pathways are depicted. Genes are represented as nodes, and biological relationship between nodes are represented as edges (lines) Genes associated with the most representative canonical pathway are highlighted in magenta.

## 4 Discussion

### 4.1. Eco-toxicological risk assessment of BP-3

The present study derived an acute toxicity value (LC50 = 3.9 mg/L) for *A. tenuis* under conditions based on OECD test guidelines. Wijgerde *et al*. (2020) performed a 6-week toxicity study of BP-3 (nominal concentration of 1 µg/L) using *A. tenuis* and indicated that BP-3 hardly affected coral survival, although the measured BP-3 concentration was drastically decreased within several days. Thus, acute adverse effects of BP-3, especially at high concentrations, on *A. tenuis* have not yet been identified. As the present study maintained the exposure concentration until the end of study, we were able to provide a reliable acute BP-3 toxicity value in *A. tenuis*.

The LC50 of BP-3 in an adult stony coral was first reported by Conway *et al*. (2021) as 4.45 mg/L (measured) in *G. fascicularis*, and the value was similar to that in the present study (3.9 mg/L), suggesting that the species sensitivity of adult corals to BP-3 was equivalent between *G. fascicularis* and *A. tenuis* (Conway et al., 2021). Moreover, the LC50 derived in this study was within the range of those of other freshwater species (ECHA, 2011; Du et al., 2017). However, because previous coral larval toxicity values were highly variable between studies (Downs et al., 2016; He et al., 2019), further studies are needed to clarify differences in BP-3 toxicity between species and life stages.

Regarding the ecotoxicological risk assessment, the PNEC was calculated as 2.0 µg/L, derived by dividing the EC50 of 2.0 mg/L in this study by the assessment factor of 1000, and the HQ was calculated as 0.67 using the highest MEC of 1.34 µg/L reported in Okinawa (Tashiro and Kameda, 2013). In addition, when using the median BP-3 concentration in a review < 0.1 µg/L (Mitchelmore et al., 2021), HQ was calculated as < 0.05. Thus, these HQ levels indicate that emission levels are properly controlled, which is corroborated by other risk assessments (Conway et al., 2021; He et al., 2019; Tsui et al., 2014). However, when outstandingly high MECs, e.g., 1.395 mg/L at Trunk Bay of St. John Island, US Virgin Islands, (Downs et al., 2016); 0.692 mg/L at a bathing spot in Galicia, Spain, (Vila et al., 2016) are used, the HQ exceeds the safety zone. The decision to use analytical values obtained in a crowded area/season and surface water should be made carefully. Moreover, the species sensitivity distribution, in which the statistical distribution of toxicity data of a chemical might be an effective approach to derive a more accurate PNEC for corals (Jung et al., 2021) and it may be desirable to integrate toxicity data from multiple species and life stages of corals.

### 4.2. Mechanistic interpretation of BP-3 toxicity and heat stress in DEG analysis and PCA

Our study provides crucial mechanistic insights into molecular mechanisms that lead to BP-3-or temperature-induced coral bleaching, and identifies potential gene markers that enable them to be distinguished. Among DEGs identified in BP-3 exposure, 78% (1,878/2,400) of the genes were BP-3 specific, indicating that *A. tenuis* responds differently to BP-3 and heat stress (Figure 2B). However, most DEGs shared by the two groups showed similar trends in expression variation, suggesting that there are responses common to both (Figure 2C).

Among shared DEGs, cytochrome P450 genes *CYP1A1* (abbreviated as *CP1A1* in the Swiss-Prot) and *CYP17A* (abbreviated as *CP17A* in Swiss-Prot) were highly upregulated by BP-3 exposure, but were suppressed by heat stress (Figure 2C). CYP1A1 is a critical enzyme in the first phase of organic pollutant catabolism (Rusni et al., 2022). Oxidation of BP-3 is catalyzed by CYP1A1 in rats (Watanabe et al., 2015) and CYP1A1 is significantly upregulated by BP-3 in *Oryzias latipes* (Kim et al., 2014) and *Danio rerio* (Meng et al., 2020), strongly suggesting the potential role of CYP1A1 in oxidative metabolism of BP-3 in *A. tenuis*. The enzyme, PGGHG, was also upregulated by BP-3, but downregulated by heat stress (Figure 2C), which is consistent with a previous study showing that PGGHG was downregulated under both heat and cold stress (Wuitchik et al., 2021). It hydrolyzes saccharide units linked to collagen in the ECM (Hamazaki and Hamazaki, 2016). The collagen α-1 (II) chain is also reportedly strongly downregulated by thermal stress (DeSalvo et al., 2010), suggesting that PGGHG expression may affect the properties of collagen. Considering that collagen is responsible for increased ECM strength and structural support of tissues, it appears that the ECM changes differently under BP-3 exposure and heat stress. Genes involved in detoxification of BP-3 including CYP1A1, and ECM-related genes such as PGGHG, are promising genetic marker candidates for determining the source of stress that affects *A. tenuis*.

To explore variation in gene expression, we performed a PCA of the identified DEGs and found that the heat-stressed group was separated by a large distance from the BP-3-exposed groups, especially by PC2 (Figure 3A), strongly suggesting differences in gene expression responses. Principal component 1 reflected the change in the concentration-dependent gene expression profile following BP-3 exposure and included the shared response between BP-3 and heat stress, which is consistent with Figures 2B and 2C. Several drug disposition-related factors, including CYP1A1, had high cos2 values and showed BP-3-specific expression changes (Figure 3B). As BP-3-specific modes of action, metabolism related changes are considered a key event. For example, MGST1, which displays both glutathione transferase and glutathione peroxidase activities and is activated by oxidative stress (Morgenstern and DePierre, 1983; Schaffert, 2011; Motone et al. 2018), may be related to detoxification of BP-3 metabolites. The MOT (Monocarboxylate transporter; MCT) gene families, which mediate cellular thyroid hormone uptake and efflux, were also identified as BP-3-specific DEGs (Figure 3B). Regulation of thyroid hormone homeostasis may be disrupted by BP-3 (Lee et al., 2018) and has a structural similarity to that of thyroxine. Further studies are needed to clarify whether MOT gene induction is involved in BP-3 transport or if it is the result of an altered hormone balance.

In addition to PGGHG (Figure 2C), ECM-related factors had high cos2 values and were highly upregulated by BP-3 exposure, suggesting that degradation and regeneration of the ECM comprises a major response to BP-3 (Figure 3C). For example, GXN is a protein component of the skeletal organic matrix (Fukuda et al., 2003) and acts as a mediator of ECM organization, oxidative stress, and defensive pathways. It was identified as a DEG in response to stony coral tissue loss disease (Traylor-Knowles et al., 2021), copper (Schwarz et al., 2013), and oil (DeLeo et al., 2018). In contrast, HMCN1 and MLP were specifically downregulated by BP-3 (Figure 3C, Table S4). HMCN is an immunoglobulin superfamily that supports architectural and structural integrity of the ECM (Xu et al., 2013) and MLP involves in synthesis of mucin. Mucin is the main component of coral mucus, which acts as a physical protective barrier. Moreover, LECG (lectin) was also substantially downregulated in BP-3_2.7 and by heat stress (Figure 3C). As certain lectins have affinities for mucin (Ahmmed et al., 2022) and express antiviral and antibacterial activity (Li et al., 2014), downregulation of lectin may reduce microbial resistance and integrity of the bacterial flora in corals. Thus, BP-3 impairs ECM homeostasis through several modes of action.

Several genes related to signal transduction-related factors were identified (Figure 3D). AGRL2 and AGRG4 are members of the GPCR. Recently, Ishii *et al*. (2022) reported that the GPCR gene expression pattern changes drastically during coral metamorphosis, suggesting that the GPCR may regulate biological and developmental processes. Furthermore, fatty acid-related enzymes (ELOV4) and development-related genes (WNT) were identified as important variables in our PCA (Figure 3D). The lipid biosynthesis enzyme, ELOV4, is differentially expressed depending on its symbiotic state (Matthews et al., 2017). Thus, alterations in fatty acid synthesis and signal transduction related to development are potentially key events in coral bleaching. Based on the combined findings of the present and previous studies, DEGs presented in this study are candidate genetic markers for coral bleaching.

A significant number of DEGs have yet to be annotated, and some of them varied remarkably in expression of BP-3 and heat-stress treatment. For example, aten_s0036.g114, aten_s0017.g163, and aten_s0024.g115 were identified as genes that change in opposite directions under heat stress and BP-3 treatment (Figure 2C). Moreover, in the PCA, aten_s0167.g11 was strongly correlated with PC2 (no correlation with PC1), indicating heat-stress-specific induction (Figure S5). Determining the molecular functions of these genes may advance our understanding of their responses to BP-3 exposure and heat stress.

### 4.3. Modes of action revealed by enrichment analyses and gene network analysis

To understand the response of *A. tenuis* to BP-3 and heat stress, GO-BP and Reactome pathway enrichment analyses were conducted for annotated genes in each group in Figure S4. The GPCR signaling pathway was the most significantly enriched GO term in BP-3-specific Up-DEGs (Figure 4), which is consistent with results shown in Figure 3D. Further studies are needed; however, GPCRs may be responsible for important BPs that occur in response to environmental stress. The “Fibroblast growth factor (FGF) receptor signaling pathway”, which was significantly enriched in BP-3-specific Up-DEGs (Figure 4), induces ECM degradation and polyp detachment via matrix metalloproteinases (MMPs) (Chuang and Mitarai, 2020). MMPs are zinc-dependent endo-peptidases, which participate in tissue remodeling during physiological and pathological processes (Hakulinen et al., 2008; Takagi et al., 2020), and several MMPs were specifically upregulated by BP-3 (Figures 3C, Table S4 and S6). Together with “extracellular matrix organization”, this strongly suggests that characteristic ECM changes occur during BP-3 exposure.

Apart from the BP-3-specific mode of action, heat-stress-specific mechanisms addressed in this study are immune system and the heat-shock response. GO terms related to the immune system were enriched in heat-specific Up-DEGs (Figure 4). An antiviral innate immune response receptor, DEAD box protein 58 (DDX58), also called as retinoic acid-inducible gene-1 (RIG-1), a member of the GO term “immune system”, were upregulated by heat stress (Table S6). DDX58 mediates innate immune response by recognizing viral nucleic acids and activating downstream signaling pathways (Matsumiya and Stafforini, 2010). Moreover, RIG-1 interacts with HSP90 and it is suggested that HSP90 protects RIG-1 from ubiquitin-mediated proteasomal degradation (Matsumiya et al., 2009). These changes could be a response to rising water temperatures, which alter the taxonomic composition of the microbiome in corals and surrounding seawater and increase the abundance of pathogenic microbes in corals (Sun et al., 2022; Thurber et al., 2009). These differences, revealed by enrichment analysis, provide an important focal point to investigate stress-specific coral bleaching mechanisms.

Pathway analysis was conducted to identify networks based on interactions between DEGs and to determine their functions. It has been hypothesized that overproduction of reactive oxygen species (ROS) in both corals and Symbiodiniaceae causes damage (Motone et al., 2020), resulting in tissue necrosis and coral cell detachment during bleaching (Gates et al., 1992). Our study suggests that multiple cellular mechanisms may also be involved in this process. Although the NRF2 pathway was identified in oxidative stress responses to both BP-3 and heat stress, the S100 family signaling pathway, which is involved in ROS induction via NADPH oxidase (NOX), was identified only in the BP-3-specific pathway (Figure 5A). Upregulated expression of tumor necrosis factor-α (TNF-α) increases ROS production by exacerbating mitochondrial dysfunction and enhancing NOX enzyme activity (Kim et al., 2007; Sumimoto et al., 2005). Since reduced nitrogen oxide (NO) levels can be related to ROS production, reduced nitric oxide synthase (NOS) might also increase ROS production (Rodrigo et al., 2011). In the present study, aten_s0007.g221 (TNF receptor-associated factor 6: TRAF6) and aten_s0012.g31 (NOX4) were increased (Figure 5A), and NOS1 was decreased after BP-3 exposure (Table S4), suggesting that these genes are related to BP-3-induced bleaching. Although NO has been proposed as an important driver of coral bleaching under heat stress (Jury et al., 2022), ROS and ONOO^-^, which can be generated by ROS and NO, may contribute to the response of corals to BP-3 exposure. Considering that MMPs contribute to the S100 signaling pathway (Figure 5A) and “extracellular matrix organization” (Table S6), and that their expression is induced by ROS (Pittayapruek et al., 2016), it is likely that the ECM state changes differently under BP-3 exposure and heat stress. Furthermore, the “G protein-coupled receptor signaling pathway” was the fourth most enriched pathway among BP-3-specific Up-DEGs, which is consistent with the GO analysis (Figure 4). Among heat-specific Up-DEGs, RIG-1 and HSP90B1 were identified in the gene network analysis (Figure 5C), suggesting heat-stress-specific interferon induction and antiviral response. Although further molecular biological studies are needed, these pathway analyses based on molecular interactions revealed both commonalities and differences in oxidative stress responses.

Consequently, an overview of differences in key events in adverse outcome pathways for BP-3-and temperature-induced bleaching can be inferred as follows. For BP-3-induced bleaching, drug metabolism and transport are crucial initiating events, and unique pathways for the ECM, e.g., FGF and MMP, and oxidative stress generating response, e.g., PDXN, NOS1, and NOX4, were identified. While this oxidative stress response was mainly caused by BP-3 exposure, subsequent NRF2-mediated oxidative stress responses were common to both BP-3 exposure and heat stress. Moreover, gene expression signatures of GPCRs and other signal transduction-related genes may be important to understand effects of stressors. For temperature-induced bleaching, immune and heat-shock responses are specific modes of action. Since many other UV filters have structural similarities to BP-3, it may be possible to distinguish between UV filters-and temperature-induced bleaching using potential gene markers identified in this study. To confirm usefulness of the gene markers, toxicity mechanisms of other UV filters on corals also need to be investigated. We hope that these differences in ecogenomic profiles can be used for environmental diagnosis to evaluate the type and intensity of stress to which each coral ecosystem is primarily subjected.

## Conclusions

The health of people, corals, and their surrounding ecosystem must be maintained through the appropriate use of UV filters. We provide reliable ecotoxicological values for BP-3 on the reef-building coral, *A. tenuis*, and show that risk is limited in most aquatic environments. To the best of our knowledge, this is the first study to report detailed differences in gene expression profiles after treatment with a UV filter (BP-3) and heat stress. These findings demonstrate differences between UV filter-and temperature-induced bleaching. The contribution of environmental stressors other than elevated sea surface temperatures in each coastal region to coral bleaching needs to be assessed. Thus, characterizing gene expression patterns of corals in each region will contribute to identifying whether the primary stressor is high water temperature or another factor such as UV filter. These diagnoses will help us realize tailor-made measures for nature-positive goals.

## Author contributions

**Sakiko Nishioka:** Formal analysis, Investigation, Methodology, Visualization, Roles/Writing – original draft. **Kaede Miyata:** Formal analysis, Investigation, Methodology, Supervision, Writing – review & editing. **Yasuaki Inoue:** Formal analysis, Investigation, Methodology, Writing – review & editing. **Kako Aoyama:** Formal analysis, Investigation, Writing – review & editing. **Yuki Yoshioka:** Formal analysis, Investigation, Methodology, Writing – review & editing. **Masayuki Yamane:** Investigation, Writing – review & editing. **Natsuko Miura:** Project administration, and Writing – review & editing. **Hiroshi Honda:** Conceptualization, Formal analysis, Investigation, Methodology, Project administration, Supervision, Visualization, Roles/Writing – original draft. **Toshiyuki Takagi:** Conceptualization, Formal analysis, Investigation, Resources, Methodology, Project administration, Supervision, Writing – review & editing.

## Acknowledgements

The authors thank Dr. Tohru Yamaguchi (Kao Corporation) for his statistical advice. Computations were partially performed on the NIG supercomputer at ROIS National Institute of Genetics.

## Research data

Raw RNA sequencing data reported are available in the DDBJ Sequenced Read Archive under the accession number ##### (to be advised prior to publication, as we are currently awaiting finalization of the data submission).

