## Supplementary materials for "Deciphering mechanisms of UV filter- and temperature-induced bleaching in the coral *Acropora tenuis*, using ecotoxicogenomics"

**Title**

**Table S1. Measured concentrations of BP-3 (mg/L) in dose range finding study (n = 1).** The retention rate at each time point compared with the initial concentration is shown in parentheses (%). LOQ: limit of quantification = 0.50 μg/L, L: low, M: medium, H: high, T: top.

| | 0 h | 24 h | 48 h | 96 h |
|---|---|---|---|---|
| BP-3_L | 0.0058 (100) | - | 0.00074 (13) | < LOQ |
| BP-3_M | 0.068 (100) | - | 0.013 (19) | 0.0055 (8.1) |
| BP-3_H | 0.69 (100) | - | 0.34 (49) | 0.18 (26) |
| BP-3_T | 6.5 (100) | 6.1 (94) | - | - |

**Table S2. Summary of water quality parameters during the 96-h treatment (n = 1).** The number after the underscore in each group name represents the time-weighted mean (TWM) concentration of BP-3. Parameters are presented as the average ± standard deviation for each test vessel. Water temperature and salinity were measured twice daily, whereas pH and dissolved oxygen (DO) were measured once a day. DO: dissolved oxygen, BC: blank control, SC: solvent control.

| | Temperature (℃) | Salinity (ppt) | pH | DO (mg/L) |
|---|---|---|---|---|
| BC | 25.9 ± 0.109 | 33.5 ± 0.108 | 8.20 ± 0.0207 | 7.80 ± 0.105 |
| SC | 25.9 ± 0.0972 | 33.6 ± 0.145 | 8.21 ± 0.0391 | 7.77 ± 0.0474 |
| BP-3_0.38 | 25.8 ± 0.109 | 33.5 ± 0.0730 | 8.20 ± 0.0297 | 7.76 ± 0.157 |
| BP-3_0.77 | 25.8 ± 0.109 | 33.5 ± 0.118 | 8.21 ± 0.0493 | 7.76 ± 0.134 |
| BP-3_1.5 | 25.8 ± 0.0866 | 33.5 ± 0.106 | 8.20 ± 0.0545 | 7.80 ± 0.0820 |
| BP-3_2.7 | 25.8 ± 0.0866 | 33.5 ± 0.0854 | 8.22 ± 0.0464 | 7.72 ± 0.0596 |
| BP-3_5.6 | 25.9 ± 0.0577 | 33.5 ± 0.0837 | 8.19 ± 0.0566 | 7.80 |
| Heat | 31.0 ± 0.0333 | 33.8 ± 0.330 | 8.22 ± 0.0919 | 6.30 ± 0.0507 |

**Table S3. Summary of read numbers and mapping rates of sequence reads.** The number after "BP-3_" in each group name represents the time-weighted mean (TWM) concentration of BP-3. The number at the end of sample details indicates the serial number of coral colony. Same number indicates that they are from the same colony. BC: blank control, SC: solvent control.

| Sample Name | Sample details | Raw read number | Trimmed read number | Mapping rate (%) |
|---|---|---|---|---|
| 1 | BC_1 | 24,157,607 | 24,128,787 | 49.79 |
| 2 | BC_2 | 27,874,431 | 27,840,954 | 45.36 |
| 3 | BC_3 | 22,637,192 | 22,612,775 | 44.23 |
| 4 | BC_4 | 26,517,965 | 26,491,222 | 52.59 |
| 5 | BC_5 | 26,407,293 | 26,376,767 | 50.15 |
| 6 | BC_6 | 26,458,600 | 26,433,628 | 49.14 |
| 7 | SC_1 | 28,582,269 | 28,548,195 | 44.86 |
| 8 | SC_2 | 25,235,334 | 25,206,117 | 43.87 |
| 9 | SC_3 | 20,979,309 | 20,957,063 | 45.62 |
| 10 | SC_4 | 26,639,280 | 26,606,934 | 50.42 |
| 11 | SC_5 | 28,921,752 | 28,885,472 | 47.90 |
| 12 | SC_6 | 24,152,082 | 24,126,414 | 49.63 |
| 13 | BP-3_0.38_1 | 20,879,341 | 20,856,932 | 44.56 |
| 14 | BP-3_0.38_2 | 23,506,354 | 23,482,562 | 41.50 |
| 15 | BP-3_0.38_3 | 21,369,600 | 21,348,782 | 44.05 |
| 16 | BP-3_0.38_4 | 26,326,990 | 26,299,860 | 50.68 |
| 17 | BP-3_0.38_5 | 20,716,342 | 20,695,558 | 47.15 |
| 18 | BP-3_0.38_6 | 32,428,047 | 32,383,474 | 48.30 |
| 19 | BP-3_0.77_1 | 24,995,587 | 24,969,263 | 43.70 |
| 20 | BP-3_0.77_2 | 28,983,434 | 28,952,988 | 44.21 |
| 21 | BP-3_0.77_3 | 27,070,836 | 27,046,089 | 45.95 |
| 22 | BP-3_0.77_4 | 24,844,283 | 24,822,825 | 52.20 |
| 23 | BP-3_0.77_5 | 22,275,220 | 22,256,693 | 46.51 |
| 24 | BP-3_0.77_6 | 22,869,155 | 22,846,857 | 47.63 |
| 25 | BP-3_1.5_1 | 27,639,358 | 27,609,868 | 47.82 |
| 26 | BP-3_1.5_2 | 29,030,125 | 28,999,526 | 45.31 |
| 27 | BP-3_1.5_3 | 30,413,583 | 30,385,238 | 51.20 |
| 28 | BP-3_1.5_4 | 21,891,468 | 21,860,874 | 48.47 |
| 29 | BP-3_1.5_5 | 23,424,408 | 23,389,502 | 49.64 |
| 30 | BP-3_1.5_6 | 22,299,954 | 22,267,615 | 50.99 |
| 31 | BP-3_2.7_1 | 21,369,912 | 21,333,162 | 41.71 |
| 32 | BP-3_2.7_2 | 26,341,822 | 26,317,276 | 43.45 |
| 33 | BP-3_2.7_3 | 22,029,237 | 22,010,792 | 46.86 |
| 34 | BP-3_2.7_4 | 24,059,445 | 24,037,517 | 52.15 |
| 35 | BP-3_2.7_5 | 23,910,277 | 23,890,422 | 46.59 |
| 36 | BP-3_2.7_6 | 24,325,216 | 24,307,601 | 51.31 |
| 37 | Heat_1 | 21,328,235 | 21,312,365 | 43.30 |
| 38 | Heat_2 | 22,176,441 | 22,156,655 | 40.60 |
| 39 | Heat_3 | 31,294,874 | 31,252,995 | 46.79 |
| 40 | Heat_4 | 22,281,009 | 22,264,811 | 48.58 |
| 41 | Heat_5 | 23,572,538 | 23,554,200 | 45.05 |
| 42 | Heat_6 | 23,080,282 | 23,059,605 | 41.91 |
| | Average | 24,888,012 | 24,861,577 | 46.95 |

**Table S4. The top 20 upregulated and downregulated differentially expressed genes (DEGs) in each gene set (Figure S4) based on log2 fold change (log2FC) compared to the control groups.** Log2FC values of BP-3_2.7 are listed as "BP-3". For common DEGs, the top 20 DEGs were extracted based on the average log2FC values of BP-3_2.7 and heat-stress groups.

| | Gene ID | UniProt ID | Discription | log2FC | |
|---|---|---|---|---|---|
| | | | | BP-3 | Heat |
| BP-3-specific Up-DEGs | aten_s0046.g96 | D9IQ16 | GXN_ACRMI_Galaxin | 9.40 | 5.74 |
| | aten_s0096.g125 | | | 8.90 | 2.04 |
| | aten_s0342.g22 | G5EG78 | PXDN2_CAEEL_Peroxidasin_homolog_pxn-2 | 8.09 | 2.32 |
| | aten_s0342.g21 | A4IGL7 | PXDN_XENTR_Peroxidasin | 7.53 | 1.03 |
| | aten_s0081.g33 | | | 6.26 | 3.12 |
| | aten_s0101.g53 | | | 6.07 | 3.03 |
| | aten_s0139.g22 | | | 5.82 | 0.81 |
| | aten_s0075.g42 | | | 5.45 | 2.24 |
| | aten_s0003.g255 | | | 5.40 | 1.61 |
| | aten_s0230.g2 | O16025 | AOSL_PLEHO_Allene_oxide_synthase-lipoxygenase_protein | 5.07 | -0.16 |
| | aten_s0085.g51 | | | 4.75 | 1.44 |
| | aten_s0012.g187 | O73853 | CP17A_ICTPU_Steroid_17-alpha-hydroxylase/17,20_lyase | 4.54 | -2.15 |
| | aten_s0016.g93 | Q90611 | MMP2_CHICK_72_kDa_type_IV_collagenase | 4.35 | 3.58 |
| | aten_s0001.g47 | Q6JZS3 | CP1A1_ORYLA_Cytochrome_P450_1A1 | 4.27 | -1.71 |
| | aten_s0005.g156 | Q8KZ94 | REBMT_LENAE_Demethylrebeccamycin-D-glucose_O-methyltransferase | 4.26 | 0.78 |
| | aten_s0079.g103 | | | 4.20 | 0.57 |
| | aten_s0005.g8 | Q32LQ4 | BHMT1_DANRE_Betaine--homocysteine_S-methyltransferase_1 | 4.07 | 0.62 |
| | aten_s0008.g170 | Q91X17 | UROM_MOUSE_Uromodulin | 4.03 | 0.90 |
| | aten_s0054.g39 | | | 3.97 | 0.12 |
| | aten_s0067.g111 | | | 3.89 | -0.47 |
| BP-3-specific Down-DEGs | aten_s0294.g5 | | | -6.61 | 0.00 |
| | aten_s0120.g1 | Q12950 | FOXD4_HUMAN_Forkhead_box_protein_D4 | -6.17 | -1.84 |
| | aten_s0026.g32 | Q60997 | DMBT1_MOUSE_Deleted_in_malignant_brain_tumors_1_protein | -6.17 | -1.61 |
| | aten_s0047.g77 | P29475 | NOS1_HUMAN_Nitric_oxide_synthase,_brain | -6.45 | -1.95 |
| | aten_s0106.g27 | P22845 | H10A_XENLA_Histone_H1.0-A | -6.41 | -0.94 |
| | aten_s0301.g2 | | | -6.28 | -1.36 |
| | aten_s0341.g15 | P84236 | H3_DROHY_Histone_H3 | -6.13 | -0.93 |
| | aten_s0143.g2 | | | -5.77 | -3.16 |
| | aten_s0069.g70 | | | -5.65 | -1.20 |
| | aten_s0117.g19 | Q9D1D6 | CTHR1_MOUSE_Collagen_triple_helix_repeat-containing_protein_1 | -5.35 | 0.12 |
| | aten_s0003.g135 | Q0V8S9 | CNTP5_CHICK_Contactin-associated_protein-like_5 | -5.31 | -0.94 |
| | aten_s0442.g3 | C0H691 | SCR2_ACRMI_Small_cysteine-rich_protein_2 | -5.18 | -1.79 |
| | aten_s0039.g46 | | | -4.86 | -0.20 |
| | aten_s0475.g5 | | | -4.96 | -1.58 |
| | aten_s0185.g26 | Q9H4G4 | GAPR1_HUMAN_Golgi-associated_plant_pathogenesis-related_protein_1 | -4.92 | -0.70 |
| | aten_s0047.g70 | | | -5.00 | -1.01 |
| | aten_s0047.g55 | | | -4.95 | -0.77 |
| | aten_s0285.g10 | Q8NDA2 | HMCN2_HUMAN_Hemicentin-2 | -4.90 | 0.42 |
| | aten_s0083.g56 | Q19319 | CADH4_CAEEL_Cadherin-4 | -4.79 | -1.17 |
| | aten_s0030.g31 | | | -4.71 | -0.86 |
| Heat-specific Up-DEGs | aten_s0029.g53 | | | -0.30 | 6.95 |
| | aten_s0091.g26 | | | 1.02 | 3.63 |
| | aten_s0093.g68 | | | 1.03 | 3.23 |
| | aten_s0051.g9 | P00973 | OAS1_HUMAN_2'-5'-oligoadenylate_synthase_1 | 1.03 | 2.94 |
| | aten_s0009.g120 | | | 0.84 | 2.85 |
| | aten_s0362.g12 | | | 0.91 | 2.75 |
| | aten_s0006.g40 | | | 1.03 | 2.61 |
| | aten_s0062.g18 | | | 0.93 | 2.34 |
| | aten_s0137.g47 | | | 1.48 | 2.34 |
| | aten_s0244.g22 | Q92058 | PPBT_CHICK_Alkaline_phosphatase,_tissue-nonspecific_isozyme | 1.38 | 2.29 |
| | aten_s0305.g9 | | | 1.02 | 2.26 |
| | aten_s0141.g3 | | | 0.92 | 2.27 |
| | aten_s0024.g115 | | | -3.01 | 2.22 |
| | aten_s0004.g79 | | | 0.50 | 2.11 |
| | aten_s0051.g10 | Q29599 | OAS1_PIG_2'-5'-oligoadenylate_synthase_1 | -0.05 | 2.05 |
| | aten_s0051.g8 | P00973 | OAS1_HUMAN_2'-5'-oligoadenylate_synthase_1 | 0.32 | 2.01 |
| | aten_s0082.g58 | B8UU74 | USOM4_ACRMI_Uncharacterized_skeletal_organic_matrix_protein_4_(Fragment) | -0.03 | 1.99 |
| | aten_s0007.g121 | D3YXG0 | HMCN1_MOUSE_Hemicentin-1 | -0.46 | 1.99 |
| | aten_s0197.g51 | | | 0.70 | 1.97 |
| | aten_s0113.g24 | | | -0.04 | 1.97 |
| Heat-specific Down-DEGs | aten_s0026.g66 | P43322 | NRG1_RAT_Pro-neuregulin-1,_membrane-bound_isoform | -5.56 | -4.50 |
| | aten_s0005.g180 | | | -2.64 | -4.45 |
| | aten_s0025.g95 | O16025 | AOSL_PLEHO_Allene_oxide_synthase-lipoxygenase_protein | -2.46 | -4.04 |
| | aten_s0007.g145 | B8V7S0 | CPP1_ACRMI_CUB_and_peptidase_domain-containing_protein_1_(Fragment) | -3.74 | -3.05 |
| | aten_s0024.g118 | P31650 | S6A11_MOUSE_Sodium-_and_chloride-dependent_GABA_transporter_3 | -1.35 | -2.92 |
| | aten_s0093.g14 | B8V7S0 | CPP1_ACRMI_CUB_and_peptidase_domain-containing_protein_1_(Fragment) | -4.61 | -2.76 |
| | aten_s0240.g27 | | | -4.23 | -2.70 |
| | aten_s0322.g13 | | | -1.41 | -2.60 |
| | aten_s0042.g26 | Q8CIW6 | S26A6_MOUSE_Solute_carrier_family_26_member_6 | -0.59 | -2.52 |
| | aten_s0032.g34 | Q5STE3 | FSTL4_MOUSE_Follistatin-related_protein_4 | -2.72 | -2.53 |
| | aten_s0031.g99 | P34329 | PDIA4_CAEEL_Probable_protein_disulfide-isomerase_A4 | -1.05 | -2.52 |
| | aten_s0007.g201 | P52569 | CTR2_HUMAN_Cationic_amino_acid_transporter_2 | -0.58 | -2.52 |
| | aten_s0005.g183 | P26445 | TNFB_PIG_Lymphotoxin-alpha | -0.56 | -2.50 |
| | aten_s0316.g13 | P05183 | CP3A2_RAT_Cytochrome_P450_3A2 | -0.49 | -2.43 |
| | aten_s0134.g23 | | | -4.42 | -2.36 |
| | aten_s0145.g6 | | | -4.15 | -2.37 |
| | aten_s0224.g11 | Q91X34 | BAAT_MOUSE_Bile_acid-CoA:amino_acid_N-acyltransferase | -2.64 | -2.36 |
| | aten_s0139.g20 | P22105 | TENX_HUMAN_Tenascin-X | -4.25 | -2.26 |
| | aten_s0014.g151 | | | -2.05 | -2.27 |
| | aten_s0029.g6 | A2BGL3 | WSCD2_DANRE_WSC_domain-containing_protein_2 | -4.15 | -2.26 |
| Common Up-DEGs | aten_s0149.g35 | P12256 | PAC_LYSSH_Penicillin_acylase | 8.41 | 7.85 |
| | aten_s0149.g36 | P12256 | PAC_LYSSH_Penicillin_acylase | 6.72 | 5.46 |
| | aten_s0117.g8 | | | 7.29 | 4.47 |
| | aten_s0004.g203 | | | 7.24 | 2.88 |
| | aten_s0114.g47 | | | 4.32 | 5.56 |
| | aten_s0051.g3 | | | 4.85 | 4.14 |
| | aten_s0197.g34 | | | 5.85 | 2.27 |
| | aten_s0095.g4 | | | 5.38 | 2.31 |
| | aten_s0056.g99 | Q6PDJ1 | CAHD1_MOUSE_VWFA_and_cache_domain-containing_protein_1 | 3.83 | 3.85 |
| | aten_s0001.g48 | | | 5.67 | 1.57 |
| | aten_s0315.g1 | | | 3.76 | 3.46 |
| | aten_s0099.g4 | | | 4.28 | 2.84 |
| | aten_s0300.g4 | | | 4.56 | 2.46 |
| | aten_s0305.g11 | | | 3.40 | 3.48 |
| | aten_s0001.g273 | O42401 | MATN3_CHICK_Matrilin-3 | 3.58 | 3.10 |
| | aten_s0285.g13 | | | 4.51 | 2.18 |
| | aten_s0313.g13 | | | 3.34 | 3.32 |
| | aten_s0576.g1 | | | 3.28 | 3.40 |
| | aten_s0010.g103 | | | 2.68 | 3.67 |
| | aten_s0235.g15 | | | 2.64 | 3.15 |
| Common Down-DEGs | aten_s0180.g46 | Q94515 | DEGS1_DROME_Sphingolipid_delta(4)-desaturase_DES1 | -10.54 | -10.45 |
| | aten_s0068.g57 | Q9EQC4 | ELOV4_MOUSE_Elongation_of_very_long_chain_fatty_acids_protein_4 | -8.13 | -7.17 |
| | aten_s0046.g80 | Q66S03 | LECG_THANI_Galactose-specific_lectin_nattectin | -10.51 | -3.65 |
| | aten_s0007.g89 | | | -8.02 | -5.17 |
| | aten_s0038.g51 | P58912 | TX60B_PHYSE_DELTA-alicitoxin-Pse2b | -8.24 | -4.48 |
| | aten_s0010.g155 | | | -9.85 | -3.14 |
| | aten_s0009.g213 | O35887 | CALU_MOUSE_Calumenin | -8.91 | -3.63 |
| | aten_s0267.g7 | | | -9.06 | -3.27 |
| | aten_s0007.g22 | B8UU51 | GXN2_ACRMI_Galaxin-2 | -8.16 | -2.74 |
| | aten_s0007.g87 | | | -6.63 | -3.29 |
| | aten_s0007.g95 | | | -5.57 | -3.93 |
| | aten_s0015.g131 | Q52SK7 | HLYS_HYDVU_Hydralysin | -4.42 | -4.23 |
| | aten_s0030.g111 | | | -5.43 | -2.99 |
| | aten_s0156.g22 | O34351 | CDLS_BACSU_Cyclo(L-leucyl-L-leucyl)_synthase | -5.96 | -2.51 |
| | aten_s0323.g13 | | | -5.48 | -3.01 |
| | aten_s0007.g86 | | | -5.19 | -3.15 |
| | aten_s0038.g49 | P58912 | TX60B_PHYSE_DELTA-alicitoxin-Pse2b | -5.35 | -3.06 |
| | aten_s0033.g40 | Q1LZE9 | PRS23_BOVIN_Serine_protease_23 | -5.59 | -2.65 |
| | aten_s0100.g7 | | | -5.70 | -2.62 |
| | aten_s0010.g7 | Q9U6Y6 | GFPL_ANEMA_GFP-like_fluorescent_chromoprotein_amFP486 | -3.20 | -4.99 |

**Table S5. Significantly enriched Gene Ontology (GO) biological process (BP) terms and Reactome pathways (FDR < 0.1) in each group.** Up-DEGs: upregulated differentially expressed genes, Down-DEGs: downregulated differentially expressed genes.

| | Category | Annotation Term | ID | No. of genes | Fold Enrichment | FDR |
|---|---|---|---|---|---|---|
| BP-3-specific Up-DEGs | GOTERM_BP_DIRECT | G-protein coupled receptor signaling pathway | GO:0007186 | 31 | 2.9595 | 0.000354 |
| | GOTERM_BP_DIRECT | extracellular matrix organization | GO:0030198 | 18 | 3.9503 | 0.002414 |
| | GOTERM_BP_DIRECT | fibroblast growth factor receptor signaling pathway | GO:0008543 | 12 | 4.1658 | 0.083327 |
| | GOTERM_BP_DIRECT | heart looping | GO:0001947 | 11 | 4.3755 | 0.089854 |
| BP-3-specific Down-DEGs | GOTERM_BP_DIRECT | chloride transmembrane transport | GO:1902476 | 14 | 5.4973 | 0.001133 |
| | GOTERM_BP_DIRECT | neurological system process | GO:0050877 | 13 | 5.7705 | 0.001133 |
| | GOTERM_BP_DIRECT | regulation of presynaptic membrane potential | GO:0099505 | 9 | 9.6720 | 0.001133 |
| | GOTERM_BP_DIRECT | chemical synaptic transmission | GO:0007268 | 20 | 3.2670 | 0.005522 |
| | GOTERM_BP_DIRECT | homophilic cell adhesion via plasma membrane adhesion molecules | GO:0007156 | 12 | 4.9005 | 0.01053 |
| | GOTERM_BP_DIRECT | excitatory postsynaptic potential | GO:0060079 | 11 | 4.9912 | 0.018915 |
| | GOTERM_BP_DIRECT | regulation of membrane potential | GO:0042391 | 13 | 3.9036 | 0.031895 |
| | GOTERM_BP_DIRECT | visual perception | GO:0007601 | 15 | 3.4031 | 0.031895 |
| | GOTERM_BP_DIRECT | neuropeptide signaling pathway | GO:0007218 | 11 | 4.4921 | 0.032309 |
| | GOTERM_BP_DIRECT | ion transmembrane transport | GO:0034220 | 14 | 3.5292 | 0.032309 |
| | GOTERM_BP_DIRECT | defense response | GO:0006952 | 8 | 5.6327 | 0.083502 |
| | GOTERM_BP_DIRECT | angiogenesis | GO:0001525 | 17 | 2.7119 | 0.095492 |
| Heat-specific Up-DEGs | REACTOME_PATHWAY | Innate Immune System | R-HSA-168249 | 7 | 6.8166 | 0.089014 |
| | REACTOME_PATHWAY | Immune System | R-HSA-168256 | 9 | 4.2118 | 0.089014 |
| Heat-specific Down-DEGs | GOTERM_BP_DIRECT | transmembrane transport | GO:0055085 | 14 | 5.3740 | 0.001465 |
| | GOTERM_BP_DIRECT | neurotransmitter transport | GO:0006836 | 5 | 16.7730 | 0.08264 |
| Common Up-DEGs | - | - | - | - | - | - |
| Common Down-DEGs | GOTERM_BP_DIRECT | transmembrane transport | GO:0055085 | 18 | 6.1084 | 3.53E-06 |
| | REACTOME_PATHWAY | SLC-mediated transmembrane transport | R-MMU-425407 | 7 | 13.0228 | 0.004102 |
| | GOTERM_BP_DIRECT | sodium ion transport | GO:0006814 | 8 | 9.7443 | 0.005952 |
| | GOTERM_BP_DIRECT | ion transport | GO:0006811 | 13 | 4.5945 | 0.006074 |
| | REACTOME_PATHWAY | Transport of small molecules | R-MMU-382551 | 9 | 5.9925 | 0.013856 |
| | REACTOME_PATHWAY | Transport of inorganic cations/anions and amino acids/oligopeptides | R-MMU-425393 | 5 | 18.6040 | 0.013856 |
| | GOTERM_BP_DIRECT | neurotransmitter transport | GO:0006836 | 5 | 14.8283 | 0.060117 |

**Table S6. Genes listed in Gene Ontology (GO) biological process (BP) terms and Reactome pathways which we focused on in the discussion.** Up-DEGs: upregulated differentially expressed genes, Down-DEGs: downregulated differentially expressed genes.

| | Category | Annotation Term | ID | UniProt ID | Gene ID | Discription |
|---|---|---|---|---|---|---|
| BP-3-specific Up-DEGs | GOTERM_BP_DIRECT | G-protein coupled receptor signaling pathway | GO:0007186 | Q9UBH6 | aten_s0053.g51 | XPR1_HUMAN_Xenotropic_and_polytropic_retrovirus_receptor_1 |
| | | | | E7F7V7 | aten_s0208.g2 | GAL2B_DANRE_Galanin_receptor_2b |
| | | | | P29274 | aten_s0197.g41 | AA2AR_HUMAN_Adenosine_receptor_A2a |
| | | | | Q96RI8 | aten_s0048.g18 | TAAR6_HUMAN_Trace_amine-associated_receptor_6 |
| | | | | O08726 | aten_s0269.g25 | GALR2_RAT_Galanin_receptor_type_2 |
| | | | | Q12802 | aten_s0033.g94 | AKP13_HUMAN_A-kinase_anchor_protein_13 |
| | | | | O93274 | aten_s0010.g125 | FZD8_XENLA_Frizzled-8 |
| | | | | Q2YDN1 | aten_s0017.g159 | GP161_BOVIN_G_protein-coupled_receptor_161 |
| | | | | Q9VCZ3 | aten_s0093.g92 | OCTB1_DROME_Octopamine_receptor_beta-1R |
| | | | | Q61212 | aten_s0099.g24 | NPY6R_MOUSE_Neuropeptide_Y_receptor_type_6 |
| | | | | Q5QD04 | aten_s0075.g100 | TAAR9_MOUSE_Trace_amine-associated_receptor_9 |
| | | | | Q96P65 | aten_s0028.g175 | QRFPR_HUMAN_Pyroglutamylated_RF-amide_peptide_receptor |
| | | | | Q18775 | aten_s1036.g2 | DOPR4_CAEEL_Dopamine_receptor_4 |
| | | | | O60242 | aten_s0060.g75 | AGRB3_HUMAN_Adhesion_G_protein-coupled_receptor_B3 |
| | | | | P0C5J4 | aten_s0041.g57 | GPR52_MOUSE_G-protein_coupled_receptor_52 |
| | | | | Q9HAR2 | aten_s0537.g1 | AGRL3_HUMAN_Adhesion_G_protein-coupled_receptor_L3 |
| | | | | G5ECQ2 | aten_s0005.g193 | FRIZ2_CAEEL_Frizzled-2 |
| | | | | Q9EQD2 | aten_s0085.g48 | NPFF2_RAT_Neuropeptide_FF_receptor_2 |
| | | | | P49219 | aten_s0003.g241 | MTR1C_XENLA_Melatonin_receptor_type_1C |
| | | | | P58308 | aten_s0042.g112 | OX2R_MOUSE_Orexin_receptor_type_2 |
| | | | | P30975 | aten_s0025.g12 | TLR2_DROME_Tachykinin-like_peptides_receptor_99D |
| | | | | Q9Y5X5 | aten_s0004.g112 | NPFF2_HUMAN_Neuropeptide_FF_receptor_2 |
| | | | | B7ZCC9 | aten_s0114.g18 | AGRG4_MOUSE_Adhesion_G-protein_coupled_receptor_G4 |
| | | | | P58421 | aten_s0055.g39 | FZD5_XENLA_Frizzled-5 |
| | | | | P35348 | aten_s0197.g42 | ADA1A_HUMAN_Alpha-1A_adrenergic_receptor |
| | | | | Q923Y3 | aten_s0038.g19 | TAA8B_RAT_Trace_amine-associated_receptor_8b |
| | | | | O57422 | aten_s0074.g7 | OPN4B_XENLA_Melanopsin-B |
| | | | | B3DM66 | aten_s0046.g17 | GP161_XENTR_G-protein_coupled_receptor_161 |
| | | | | Q8ITC7 | aten_s0052.g81 | CAPAR_DROME_Neuropeptides_capa_receptor |
| | | | | C3ZQF9 | aten_s0030.g19 | QRFPR_BRAFL_QRFP-like_peptide_receptor |
| | | | | P47900 | aten_s0058.g18 | P2RY1_HUMAN_P2Y_purinoceptor_1 |
| | GOTERM_BP_DIRECT | extracellular matrix organization | GO:0030198 | Q9UKP4 | aten_s0018.g60 | ATS7_HUMAN_A_disintegrin_and_metalloproteinase_with_thrombospondin_motifs_7 |
| | | | | Q9Y5R2 | aten_s0180.g67 | MMP24_HUMAN_Matrix_metalloproteinase-24 |
| | | | | Q8C6K9 | aten_s0215.g8 | CO6A6_MOUSE_Collagen_alpha-6(VI)_chain |
| | | | | Q9ER58 | aten_s0115.g14 | TICN2_MOUSE_Testican-2 |
| | | | | P59511 | aten_s0056.g40 | ATS20_MOUSE_A_disintegrin_and_metalloproteinase_with_thrombospondin_motifs_20 |
| | | | | Q3UQ28 | aten_s0004.g169 | PXDN_MOUSE_Peroxidasin_homolog |
| | | | | Q3U435 | aten_s0180.g62 | MMP25_MOUSE_Matrix_metalloproteinase-25 |
| | | | | Q8TE57 | aten_s0003.g248 | ATS16_HUMAN_A_disintegrin_and_metalloproteinase_with_thrombospondin_motifs_16 |
| | | | | Q868Z9 | aten_s0045.g7 | PPN_DROME_Papilin |
| | | | | Q9P2N4 | aten_s0056.g39 | ATS9_HUMAN_A_disintegrin_and_metalloproteinase_with_thrombospondin_motifs_9 |
| | | | | Q68BL8 | aten_s0018.g38 | OLM2B_HUMAN_Olfactomedin-like_protein_2B |
| | | | | P21359 | aten_s0001.g66 | NF1_HUMAN_Neurofibromin |
| | | | | Q99542 | aten_s0061.g38 | MMP19_HUMAN_Matrix_metalloproteinase-19 |
| | | | | P08120 | aten_s0027.g89 | CO4A1_DROME_Collagen_alpha-1(IV)_chain |
| | | | | A6NMZ7 | aten_s0215.g5 | CO6A6_HUMAN_Collagen_alpha-6(VI)_chain |
| | | | | A6H584 | aten_s0016.g64 | CO6A5_MOUSE_Collagen_alpha-5(VI)_chain |
| | | | | Q90611 | aten_s0016.g93 | MMP2_CHICK_72_kDa_type_IV_collagenase |
| | | | | P22105 | aten_s0044.g130 | TENX_HUMAN_Tenascin-X |
| | GOTERM_BP_DIRECT | fibroblast growth factor receptor signaling pathway | GO:0008543 | Q0IH87 | aten_s0120.g5 | PAX3B_XENLA_Paired_box_protein_Pax-3-B |
| | | | | P48804 | aten_s0111.g23 | FGF4_CHICK_Fibroblast_growth_factor_4 |
| | | | | E7FAM5 | aten_s0035.g111 | LIN41_DANRE_E3_ubiquitin-protein_ligase_TRIM71 |
| | | | | Q6V0I7 | aten_s0072.g28 | FAT4_HUMAN_Protocadherin_Fat_4 |
| | | | | Q498D6 | aten_s0076.g44 | FGFR4_RAT_Fibroblast_growth_factor_receptor_4 |
| | | | | P22607 | aten_s0408.g2 | FGFR3_HUMAN_Fibroblast_growth_factor_receptor_3 |
| | | | | Q9NP95 | aten_s0170.g48 | FGF20_HUMAN_Fibroblast_growth_factor_20 |
| | | | | Q7SIF8 | aten_s0015.g81 | FGF1_NOTVI_Fibroblast_growth_factor_1_(Fragment) |
| | | | | Q03364 | aten_s0287.g16 | FGFR2_XENLA_Fibroblast_growth_factor_receptor_2 |
| | | | | P34004 | aten_s0168.g8 | FGF1_MESAU_Fibroblast_growth_factor_1 |
| | | | | Q2PZL6 | aten_s0001.g171 | FAT4_MOUSE_Protocadherin_Fat_4 |
| | | | | Q6I6M7 | aten_s0115.g41 | FGF1_CYNPY_Fibroblast_growth_factor_1_(Fragment) |
| Heat-specific Up-DEGs | REACTOME_PATHWAY | Immune System | R-HSA-168256 | O95786 | aten_s0189.g41 | DDX58_HUMAN_Antiviral_innate_immune_response_receptor_RIG-I |
| | | | | P10914 | aten_s0082.g67 | IRF1_HUMAN_Interferon_regulatory_factor_1 |
| | | | | Q9Y2Z0 | aten_s0136.g37 | SGT1_HUMAN_Protein_SGT1_homolog |
| | | | | P00519 | aten_s0019.g46 | ABL1_HUMAN_Tyrosine-protein_kinase_ABL1 |
| | | | | P21580 | aten_s0095.g37 | TNAP3_HUMAN_Tumor_necrosis_factor_alpha-induced_protein_3 |
| | | | | P00973 | aten_s0051.g8 | OAS1_HUMAN_2'-5'-oligoadenylate_synthase_1 |
| | | | | Q13233 | aten_s0079.g110 | M3K1_HUMAN_Mitogen-activated_protein_kinase_kinase_kinase_1 |
| | | | | Q15399 | aten_s0135.g69 | TLR1_HUMAN_Toll-like_receptor_1 |
| | | | | Q7RTR2 | aten_s0213.g8 | NLRC3_HUMAN_NLR_family_CARD_domain-containing_protein_3 |

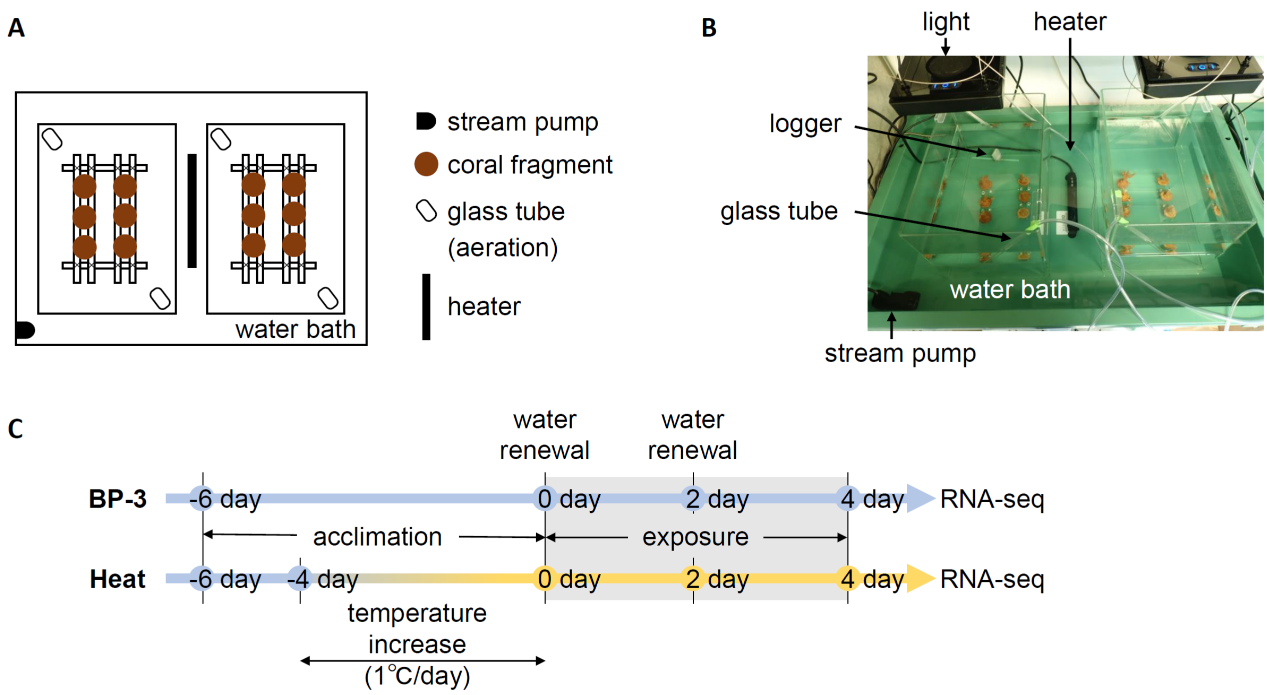


**Figure S1. Overview of the experimental design.** (A, B) Overview of experimental setup. Two glass vessels were set in a water bath, and water temperature of the water bath was controlled using a heater and a stream pump. A vessel for heat-stress experiment was set in a separate water bath. Six fragments from different colonies were placed in each test vessel (n = 6). Two glass tubes were used for aeration in each vessel to maintain dissolved oxygen (DO) levels and to provide constant random water flow. (C) A graphical summary of the experimental timeline. Upper row shows the acute-toxicity test (96 h) timeline for the control (BC, SC) and BP-3 exposure groups, and lower row shows that for the heat-stress group. Coral fragments were acclimatized for six days immediately before the start of exposure, and complete renewal of solutions was carried out at the start of exposure and at 48 h (semi-static). Water temperature of the heat-stress group was increased from 26 to 31° C at a rate of 1° C per day after acclimatization at 26° C for two days.

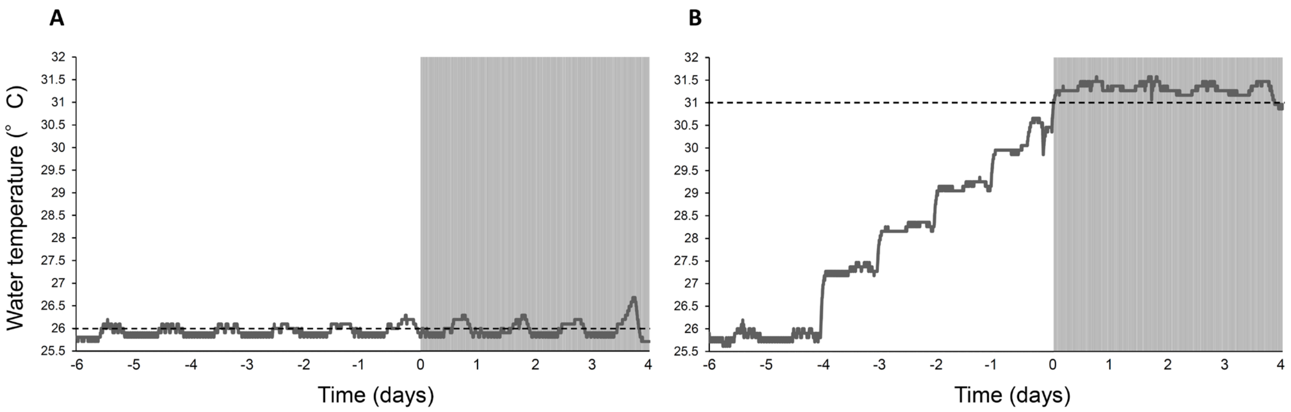


**Figure S2. Water temperatures of blank control group (A) and heat-stress group (B) throughout the experiment.** They were recorded every 5 min throughout the experiment. For the heat-stress group, water temperature was increased from 26 to 31° C at a rate of 1° C per day after two days of acclimatization at 26 ° C. The gray shading shows the exposure period, and dotted line shows the set temperatures of each group.

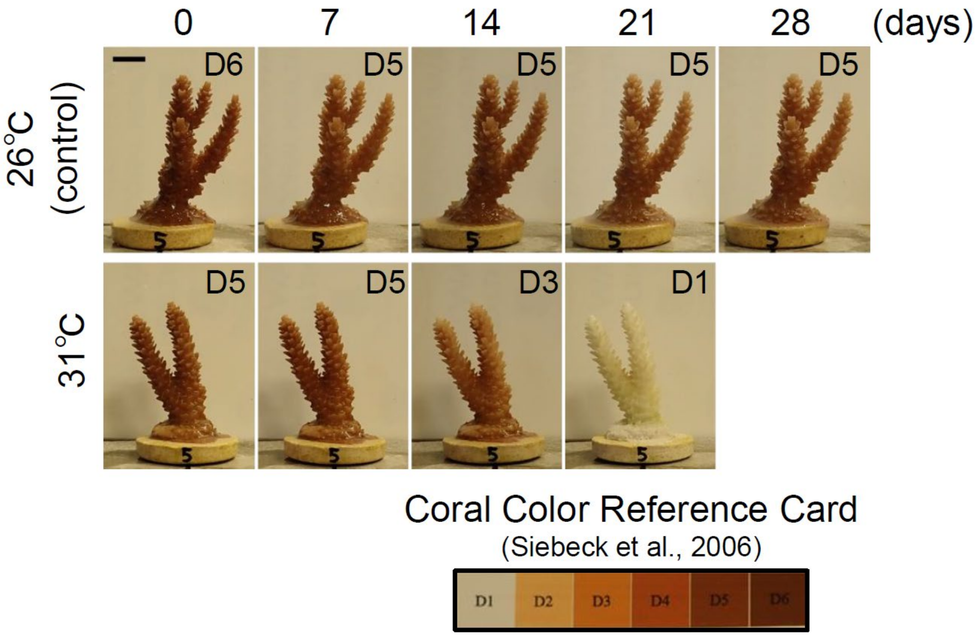


**Figure S3. Long-term exposure to heat stress.** *Acropora tenuis* fragments from the same colonies as in the BP-3 exposure test were used. Water temperature was increased from 26 to 31° C at a rate of 1° C per day, and the fragment was exposed to 31° C for 28 days (4 weeks) after reaching 31° C. Coral health was maintained even after 28 days in control group (26° C). Bleaching became gradually conspicuous from the 14th day in 31° C group, and fragments were completely bleached on the 21st day, confirming that this test system could reproduce coral bleaching phenomenon. Numbers in the upper right corner of each photo correspond to scores of the coral color reference card (Siebeck et al., 2006). Bar = 1 cm.

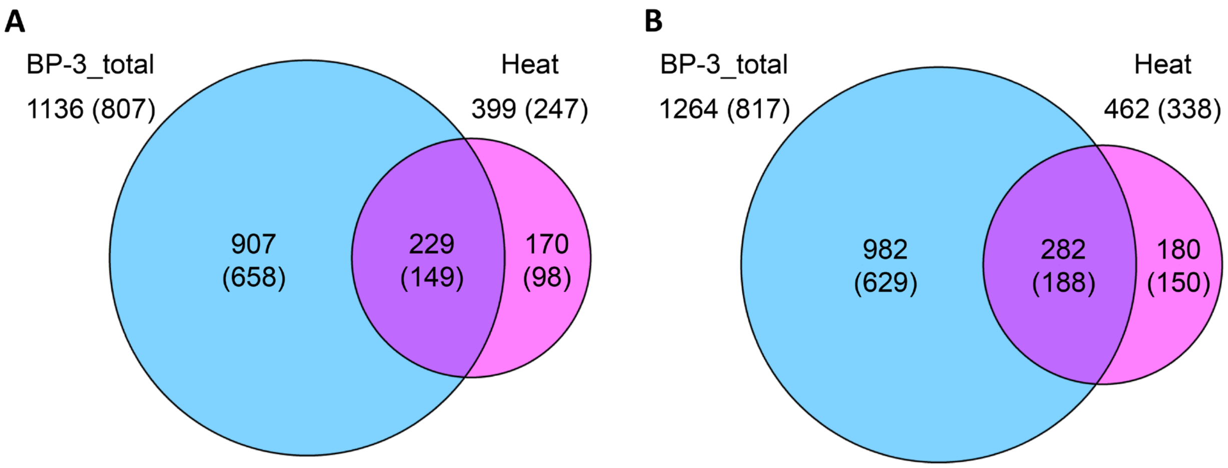


**Figure S4. Comparison of differentially expressed genes (DEGs) between BP-3 (cyan) and heat-stress exposure (magenta) groups.** Venn diagrams of numbers of upregulated DEGs (A) and downregulated DEGs (B) showing expression variation upon exposure to any concentration of BP-3 (BP-3_total) and DEGs upon heat stress (Heat). Numbers of annotated DEGs in Swiss-Prot database are shown in parentheses.

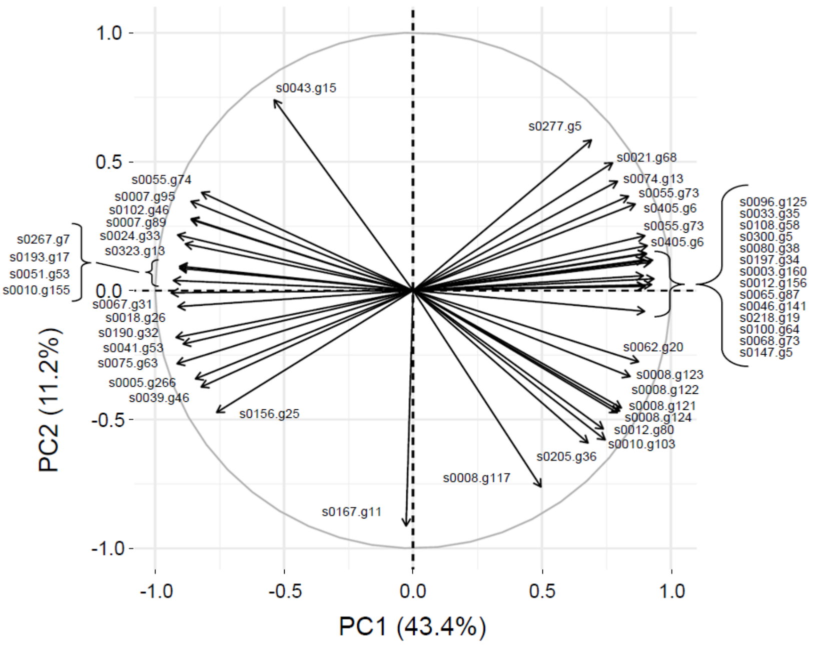


**Figure S5. Principal component analysis (PCA) biplot extracted for unannotated differentially expressed genes (DEGs) with cos2 > 0.8.** PCA biplot is based on log2FC values of all DEGs that showed expression differences in control and treatment groups (Figure 3A). Contributions of the first (PC1) and second principal component (PC2) were 43.4% and 11.2%, respectively. The initial "aten_" is omitted for each gene name.
